# Next-generation enhancer AAVs for selective interspecies targeting of midbrain dopaminergic neurons

**DOI:** 10.1101/2025.11.06.687071

**Authors:** Kian A. Caplan, Ke Chen, Martin Wienisch, Yongqi Wang, Ruoyang Chai, In-Hye Kang, Cindy Szyin Chen, Ricardo del Rosario, Fenna M. Krienen, Fan Wang, Guoping Feng

**Author notes:** Authors contributed equally (Ke Chen, Martin Wienisch, Yongqi Wang).

## Abstract

Using single-nucleus multiomic profiling of the marmoset midbrain, we develop and benchmark enhancer–AAVs to selectively access dopamine (DA) neurons in wild-type animals without combinatorial systems. *In vivo* candidate screenings identified one highly specific DA enhancer, cjDAE8, in both mice and marmosets. To overcome low-level off-target (leaky) expression, a common limitation for enhancer AAVs, we engineered next-generation AAV backbones that strengthen expression while minimizing leakiness. Quantitative histology comparing natural versus antibody-amplified fluorescence defined AAV doses to achieve high labeling efficiency with greater than 90–95% DA-neuron specificity across species and injection routes. We further demonstrate applications of DA-enhancer-AAVs for (i) retrograde targeting of projection-defined DA populations in marmoset, (ii) fiber-photometric recording of divergent DA-axonal dynamics in striatal subregions, and (iii) optogenetic VTA-DA self-stimulation in mice. Our results establish a resource for cross-species DA targeting and two practical guidelines: backbone context critically shapes enhancer performance, and antibody-amplified readouts rigorously assess specificity.

## Introduction

Dopamine (DA) plays a critical role in motor control, motivation, reward processing, and learning. Unsurprisingly, DA neuron dysfunction is featured across the clinical landscape, with disorders ranging from neurological (e.g., Parkinson’s disease)^1–5^ to psychiatric (e.g., ADHD, addiction, schizophrenia)^6–10^. The projections of DA neurons are widespread and far-reaching, with distinct projection targets and functional specialization between terminals arising from the substantia nigra pars compacta (SNc) and ventral tegmental area (VTA)^11^. Canonically, SNc DA neurons regulate voluntary movements and motor learning through axon terminals that densely innervate the basal ganglia, forming the nigrostriatal pathway^1,2,12–14^. In contrast, VTA DA neurons are crucial for motivated behavior and reward learning through their dense innervation of the nucleus accumbens, cortex, and other limbic regions, forming the mesocorticolimbic pathway^15–17^. To selectively access DA neurons in rodents, DA transporter (DAT)-IRES-Cre mice are widely used^18,19^. Among pan tyrosine hydroxylase (TH)-Cre and DAT-Cre lines, only DAT-IRES-Cre mice achieved >90% specificity of transgene expression in the literature, with others ranging between ∼40–70%, when crossed with Cre-dependent reporter lines^20,21^ or injected with Cre-dependent AAV expression cassettes^22–27^. While DAT-Cre mice are an invaluable research tool, there are still caveats to their use. DAT protein levels are reduced by nearly 17% and 50% in heterozygotes and homozygotes, respectively^28^. Furthermore, off-target Cre expression in nondopaminergic populations of limbic brain regions (amygdala, septum, hypothalamus, habenula)^29^ and abnormal behaviors (baseline hyperactivity; reduced locomotion/stereotypy after amphetamine)^30^ are reported.

These features raise concerns that the DAT-Cre knock-in may perturb cellular activity and circuit dynamics. Thus, even in rodents, there is a need for tools to selectively target DA neurons without disturbing native transcriptional programs, circuit function, or behavior. In non-human primates (NHPs), selective DA-neuron targeting has previously been pursued using transgenic and viral strategies, with varying success. Yoshimatsu et al. (2023) characterized the first founder TH-Cre marmoset but reports on the selectivity and efficiency of Cre-dependent targeting await offspring^31^, and the ∼1.5-year sexual maturation and breeding cost of marmosets limits its widespread availability^32^ and utility for studying DA system dysfunction in other genetically engineered marmoset disease models. Non-transgenic approaches include intersectional viral strategies based on connectivity (e.g., retrograde AAV-Cre in a downstream target combined with Cre-dependent AAV in midbrain) and cell-type specific promoters (e.g., AAV driving Cre expression under control of the TH promoter or lentivirus directly driving transgene under control of the TH promoter)^33–36;^ each faces challenges with specificity/efficiency, reproducibility, or payload size. Collectively, the field still lacks fast-acting and physiologically faithful tools to access NHP DA neurons with high specificity and efficiency.

In recent years, enhancer AAVs have emerged as a promising tool for selective genetic access to distinct brain cell types, with varying granularity (i.e., region- and subtype-selectivity). Nominated candidate sequences are cloned upstream of a minimal promoter in modified AAV constructs and screened (*in vitro* or *in vivo*) to identify enhancers driving robust, efficient, and specific expression in targeted cell types^27,37–44^. Given that enhancer activity can have remarkable evolutionary conservation, enhancer AAVs are uniquely positioned for selective cell-type targeting across species. Importantly, the technology can either circumvent transgenic animal lines altogether or be used in combination for multiplexed targeting and intersectional strategies. Enhancer AAVs have been reported for interspecies (e.g., rodent, NHP, avian) targeting of cortical and subcortical interneuron populations^37–41,43–47^, layer-specific cortical excitatory projection neurons^38,43,45^, striatal interneurons and projection neurons^37,48,49^, as well as granule cells of the dentate gyrus^40^. Many of these cited enhancer tools are featured in the Genetic Tools Atlas (RRID: SCR_025643), a publicly available resource by the Allen Institute that screened 1,720 enhancer AAV candidates for 44 different neuron and glial cell types. The Genetic Tools Atlas screened 95 enhancer candidates for midbrain dopamine (DA) neurons in mouse, reporting one candidate (AiE0888m) with high efficiency and >90% specificity (unpublished AiP13863, Addgene #199773). Yet, to our knowledge, no enhancer AAVs have been reported that specifically target midbrain DA neurons in NHPs.

Here, we developed an enhancer-AAV toolbox to label, record, and manipulate DA neurons, or projection-defined DA neurons, across species. Regardless of local or systemic delivery, we achieved unprecedented specificity and efficiency in mice and marmosets by using modified, next-generation enhancer-AAV backbone vectors to augment the strength and minimize leakiness of transgene expression. Through a combination of immunofluorescent staining, imaging in serially sectioned or cleared brain samples (epifluorescent, confocal, or light-sheet), and semi-automated histology analysis pipelines for robust and comprehensive quantification, we characterized AAV doses for local and systemic administration that achieve high efficiency labeling with >90% specificity in mouse and >95% specificity in marmoset. To demonstrate the utility of cjDAE AAVs as a powerful *in vivo* research tool for DA neurons, we i) labeled projection-target defined DA neurons in the marmoset using a retro-AAV approach, ii) expressed GCaMP6f in the SNc and recorded highly divergent calcium transients between DA terminals in the central striatum (CS) and dorsolateral striatum (DLS) across behavioral contexts in mice using fiber photometry, and iii) expressed channelrhodopsin in the VTA and elicited robust nose-poke triggered self-stimulation of DA neurons in mice. Taken together, our validated cjDAE AAVs provide a more accessible alternative approach for selectively targeting DA neurons in rodents and NHPs with comparable specificity and efficiency to gold-standard approaches.

## Results

### Nomination of cjDAE candidates and preliminary *in vivo* mouse screen

Dual 10X snRNA- and snATAC-sequencing (2x reactions) of unsorted nuclei isolated from marmoset ventral midbrain tissue generated 32,070 high-quality nuclei for integrated multiomic analyses (Extended Data Fig. 1a, see methods for details). A total of 258 DA nuclei across combined reaction datasets were used to identify differentially accessible chromatin peaks. We restricted the list of putative DA-neuron enhancer peaks to 401 candidates by thresholding based on the specificity of peak penetrance (>30% in target cluster, <5% in off-target) and normalized magnitude of peak expression (read counts ≥1.5 log2 fold-change) in DA neurons compared to non-DA clusters (Extended Data Fig. 1b). Among candidate peaks with the highest penetrance and expression levels in DA neurons, we further considered genes proximal to the peak in linear sequence distance and their mRNA enrichment in DA neurons (Extended Data Fig. 1c-d), together with cross-species sequence conservation (Extended Data Fig. 2) and genome context, to select 12 candidates for *in vivo* screening. For example, the cjDAE8 peak was highly specific (41.9% DA vs 3.8% non-DA penetrance) and nearly 800 bp, located in the intron of LOC100413948, an uncharacterized protein-coding locus. Although LOC100413948 is expressed by all DA neurons at high levels, it is nonspecific to this population (Extended Data Fig. 1c). In addition, the cjDAE4 peak was also highly specific (31.8% DA vs 1.4% non-DA penetrance), nearly 900 bp, and in the intron of another uncharacterized locus, LOC118155346 (Extended Data Fig. 1d). However, unlike LOC100413948 for cjDAE8, the expression of this locus is highly downregulated in all DA neurons. Regarding evolutionary conservation, both cjDAE8 (Extended Data Fig. 2a) and cjDAE4 (Extended Data Fig. 2b) contained highly conserved DNA regions across primates and mammals. Together, these criteria were considered to select the first 12 candidates for *in vivo* screening.

To screen for putative DA enhancers driving DA-specific *in vivo* expression, cjDAE1-cjDAE12 peaks were cloned into conventional enhancer pAAV vectors described previously^43,49^ expressing either EGFP or dTomato (dTom) to screen candidates in pairs. Between AAV ITR sequences, the expression cassette contained an inverted Woodchuck hepatitis virus Post-transcriptional Regulatory Element (iWPRE) as an insulator, the amplified enhancer sequence, a cytomegalovirus (CMV) minimal promoter (CMVmini2), EGFP (cjDAE1-E6) or dTomato (cjDAE7-E12) reporter, WPRE, and a modified bovine growth hormone poly A (mBGHpA). We used a non-titered, minimal purification approach^50^ after packaging the constructs in AAV-PHP.eB for the initial screening to maximize efficiency. Mice (n=6) were systemically injected (via retro-orbital route) with EGFP- and dTomato-expressing candidate pairs to qualitatively assess the relative specificity and efficiency of reporter expression in TH+ DA neurons (Fig. 1a). Out of all 12 cjDAE candidates, only six drove expression in DA neurons. Candidates not selected either failed to express the reporter in DA neurons, with some sparse labeling throughout the brain, if at all (n=2), or showed robust off-target expression of the reporter (n=4). The six candidates with discernible reporter expression in DA neurons varied widely in their specificity and labeling efficiency. For two candidates, cjDAE8-dTomato and cjDAE4-EGFP, we observed strikingly robust and highly specific reporter expression in midbrain DA neurons (Fig. 1b) with minimal off-target labeling throughout the brain. We therefore selected these two candidates for *in vivo* testing in the marmoset.

**Figure 1.**
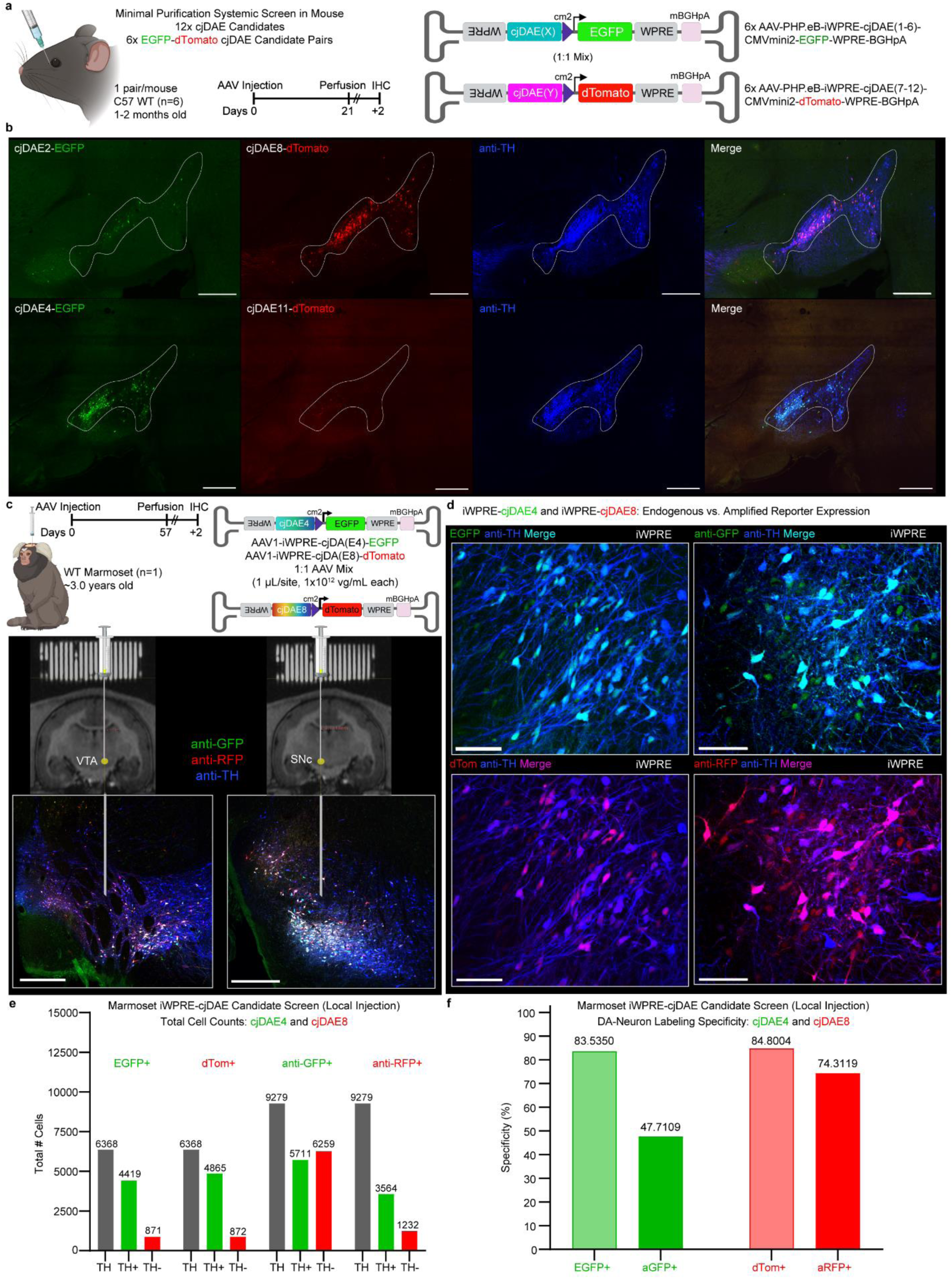
Mouse and marmoset *in vivo* screen of cjDAE-AAVs identifies two lead candidates, cjDAE8 and cjDAE4. **a,** Enhancer candidate peaks, amplified from marmoset gDNA, were cloned into pAAV constructs expressing either EGFP or dTomato and screened in mice (n=6, 2-3 months old) by systemic injection with minimally purified AAV-PHP.eB preparations of EGFP-dTomato candidate pairs. **b,** Only two candidates, cjDAE4 and cjDAE8, exhibited robust and specific reporter expression and were selected for screening in a marmoset. Scale bar, 500 μm. Schematics made in BioRender. **c,** Screening strategy and timeline for local injection of cjDAE4-EGFP and cjDAE8-dTomato in marmoset using a conventional enhancer-AAV backbone (iWPRE-insulator, mBGHpA). Stereotactic intraparenchymal AAV injections (1:1 mix, 1E+12 vg/ml each) were delivered to the marmoset SNc and VTA by combined implantation of coordinate-grid and structural MRI imaging. Representative images of the injection sites show restricted transgene expression in both regions. Scale bar, 500 μm. **d,** Representative confocal images of endogenous (EGFP/dTom) versus amplified (aGFP/aRFP) reporter expression at the injection site for cjDAE4 and cjDAE8. Scale bar, 100 μm. **e,** Quantification of aggregated cell counts for endogenous reporter expression surrounding the injection site versus amplified reporter expression sampling the entire midbrain. **f,** Summary of specificity for endogenous versus amplified DA-neuron labeling of cjDAE4 (EGFP+/aGFP+) and cjDAE8 (dTom+/aRFP+). As the most specific candidate after antibody amplification (74.31% versus 47.71%, cjDAE8 was selected for further characterization and optimization. ***Definitions*.** Specificity = (reporter⁺TH⁺ / reporter⁺ total)×100; Efficiency = (reporter⁺TH⁺ / TH⁺ total)×100. Marmoset and AAV schematics made in BioRender.

### *In vivo* test of cjDAE4 and cjDAE8 in marmoset

cjDAE4-EGFP and cjDAE8-dTomato constructs, in their original form, were packaged in AAV1, mixed 1:1, and co-injected into the SNc and VTA (1E+12 vg/ml each, one μl/site) of one male marmoset (aged ∼3 years) through MRI-targeted stereotactic surgery. After 59 days of incubation, both cjDAE4-EGFP and cjDAE8-dTomato AAVs engendered relatively dim but significant reporter expression in DA neurons (Fig. 1c). Because the spread of reporter expression was limited, we assessed native reporter expression (endogenous fluorescence without antibody amplification) using serial coronal sections of the apex of the injection site. After immunostaining to label DA neurons by their expression of the tyrosine hydroxylase (TH) protein, we quantified 6,368 anti-TH+ cells, 5,290 EGFP+ cells, and 5,737 dTom+ cells total across sections. With 871 EGFP+TH-(off target) cells for cjDAE4, reporter labeling was 83.54% specific to DA neurons (Fig. 1d-f). Similarly, for cjDAE8, we detected 4865 dTom+TH+ (on target) cells and 872 dTom+TH-(off target) cells, resulting in 84.80% specific labeling (Fig. 1d-f). Next, we amplified reporter expression by immunostaining, using serial sections (300 μm apart) spanning the midbrain, with at least one representative section from each injection site. In total, we quantified 9,279 total anti-TH+ cells, 11,970 anti-GFP+ cells, and 4,796 anti-RFP+ cells. The specificity of cjDAE4 dramatically reduced to 47.71%, with 6,259 anti-GFP+TH-cells detected. In contrast, for cjDAE8, only 1,232 anti-RFP+TH-cells were observed, maintaining a higher specificity of 74.31% (Fig. 1d-f) compared to cjDAE4. We therefore selected cjDAE8 for further validation in marmosets and mice.

Although we are confident in the accuracy of cell counts for antibody-amplified cjDAE4 reporter expression, the naturally high autofluorescence in the 488-nm channel, combined with dim transgene expression in the unamplified dataset, proved challenging for automated detection of dim off-target EGFP+ cells while also sparing false-positive detections of background fluorescence. We therefore used a more conservative intensity threshold for the unamplified cjDAE4 channel, which artificially increased specificity by 10-20% relative to manual counts.

To achieve optimal and tight transgene expression across administration routes for both species, we inserted cjDAE8 into modified enhancer AAV backbones featuring a tandem chicken hypersensitive site 4 (2xCHS4)^51–54^ insulating sequence to mitigate ITR-enhancer interactions, as well as a truncated human growth hormone polyA (mHGHpA) to improve expression strength in the marmoset.

### Pan and projection-defined targeting of DA neurons in the marmoset by local delivery of optimized cjDAE8-AAVs

Using the same AAV1 serotype, dosage, incubation time, and fluorescent reporter (dTom), one male marmoset ( aged 2.32 years) received stereotactic injections (1E+12 vg/ml, one μl/site) at two midbrain sites with the modified 2xCHS4-cjDAE8-dTom AAV vector to directly assess whether modifications to AAV backbone altered the specificity and efficiency of reporter expression for the same enhancer (Fig 2a; Extended Data Fig. 3). The multiplicity of transcriptionally distinct DA-neuron subtypes engenders diverging patterns of axon terminal projections with complex functional specialization^3,55^. Therefore, we packaged the same construct, driving EGFP expression, in the AAV2retro serotype and unilaterally injected the posterior caudate and putamen (5E+12 vg/ml, 1.5 μl/site) to further stratify DA neuron labeling by projection targets (Fig. 2a; Extended Data Fig. 3).

**Figure 2.**
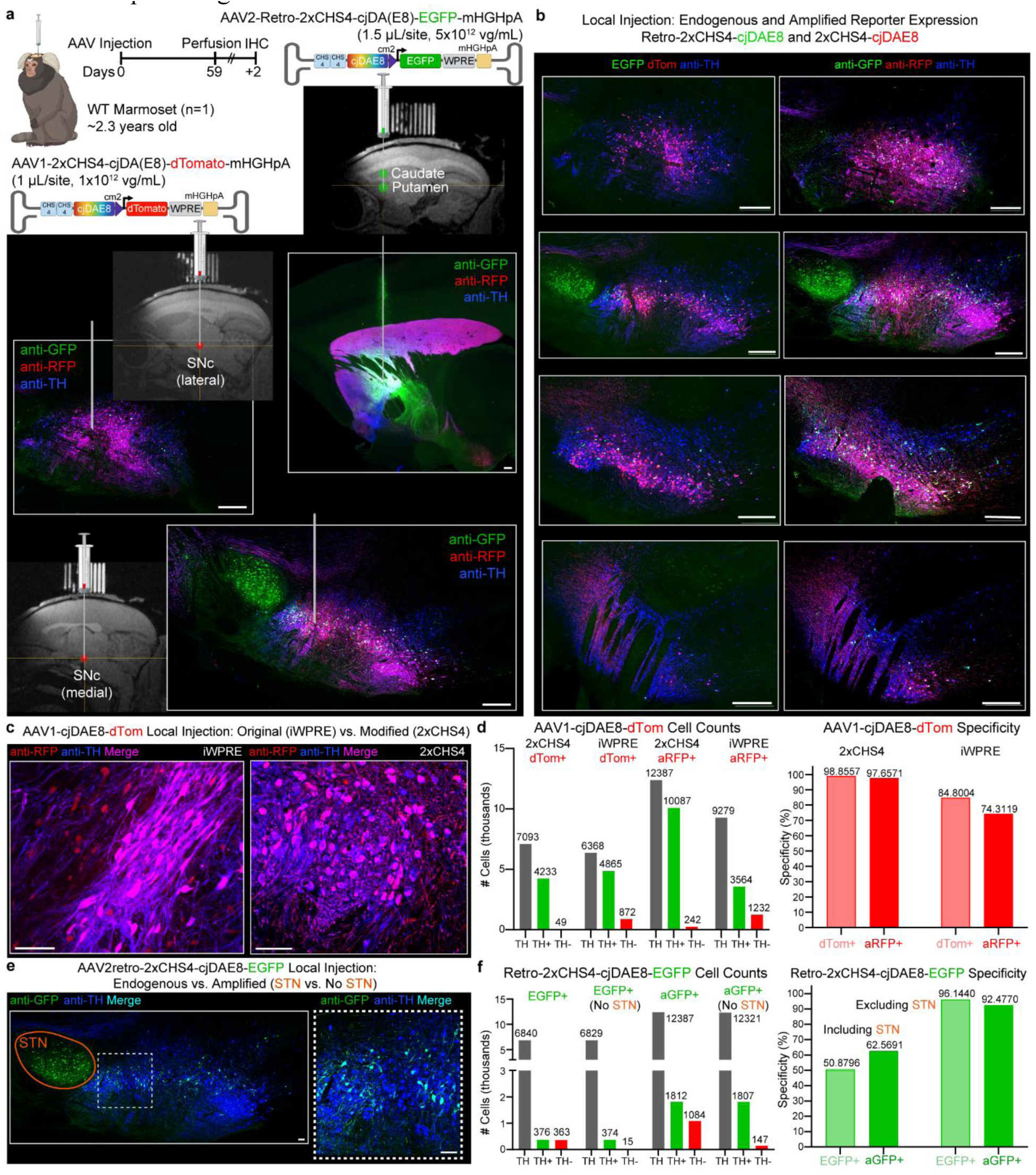
Local injection of backbone-modified cjDAE8 AAVs in the marmoset enables robust, selective transgene delivery in pan or projection-defined DA neurons. **a,** Experimental strategy and timeline for local injection of backbone-modified cjDAE8 AAVs in marmoset to achieve direct or retrograde targeting of DA. To enable comparisons with the screening dataset, identical or near identical marmoset age, AAV capsid, dose, incubation time, and reporter fluorophore were used for the 2xCHS4-cjDAE8-dTom construct. In contrast, AAV2-retro-2xCHS4-cjDAE8-EGFP was injected into the caudate and putamen to label DA neurons by projection-target. Representative images of MRI-targeted stereotaxic injections in the caudate and putamen (Retro-EGFP; 5E+12 vg/mL, 1.5 μL/site), as well as the lateral and medial SNc (dTom; 1E+12 vg/mL, 1 μL/site). Scale bar, 500 μm. **b,** Representative confocal images of endogenous (dTom/EGFP) versus amplified (aRFP/aGFP) reporter expression across lateral-medial axis of the midbrain. Scale bar, 500 μm. Marmoset and AAV schematics made in BioRender. **c,** Representative confocal images of antibody-amplified reporter expression at the injection site for cjDAE8 with the original (iWPRE-insulator) versus modified (2xCHS4-insulator) enhancer AAV backbones. Scale bar, 100 μm. **d,** Quantification of aggregated cell counts and specificity for endogenous versus amplified dTomato expression between backbones. Despite equivalent AAV capsids, dosing, reporter, and incubation time, the modified 2xCHS4-cjDAE8 construct dramatically improved the specificity of both endogenous and antibody-amplified reporter labeling compared with iWPRE-cjDAE8. **e,** Representative confocal images of antibody-amplified retrograde transduction of midbrain DA neurons projecting to subregions of the caudate and putamen. Scale bar, 100 μm. **f,** Posterior spread of the AAV injection into the globus pallidus caused robust transduction of the subthalamic nucleus (STN). However, within the midbrain, both endogenous and antibody-amplified reporter expression surpassed 90% specificity in DA neurons. ***Definitions*.** Specificity = (reporter⁺TH⁺ / reporter⁺ total)×100; Efficiency = (reporter⁺TH⁺ / TH⁺ total)×100.

Remarkably, local injection of the modified 2xCHS4-cjDAE8-dTom AAV dramatically improved the strength and specificity of transgene expression (Fig. 2b). For the endogenous-fluorescence dataset, we sampled the entire midbrain using serial sagittal sections spaced 300 μm apart. In total, we quantified 7,093 anti-TH+ cells and 4,282 dTom+ cells. With only 49 dTom+TH-(off-target) cells detected, the specificity of cjDAE8 labeling in DA neurons improved to 98.86% (Figs. 2b-d). For the reporter-amplified dataset, we further subsampled the midbrain at 100-μm intervals and quantified 12,387 anti-TH+ cells and 10,329 anti-RFP+ cells total. Remarkably, reporter labeling retained 97.66% specificity after antibody amplification, with only 242 anti-RFP+TH-cells detected (Fig. 2b-d). Thus, compared with the conventional iWPRE-cjDAE8 AAV, local injection of 2xCHS4-cjDAE8-dTom AAV improved the specificity of endogenous reporter labeling and remained unchanged after antibody amplification (Fig. 2c-d; Extended Data Fig. 3). Taken together, these data suggest i) that local injection of cjDAE8 AAV is a high-performance method for targeting midbrain DA neurons in the marmoset, and also that ii) elements of the enhancer-AAV construct backbone sequence (e.g., insulator, polyA) significantly contribute to the strength and specificity of transgene expression.

Despite conflicting reports regarding the transduction efficiency of AAV2retro in NHP DA neurons with conventional promoter-based expression cassettes^56,57^, we observed robust AAV2retro-2xCHS4-cjDAE8-EGFP labeling in spatially localized midbrain DA-neuron populations for both native and antibody-amplified expression (Fig. 2e-f; Extemded Data Fig 3b-c). Unamplified fluorescence showed specific retrograde EGFP labeling of midbrain DA neurons across brain sections, with no labeling in other canonical striatal-projecting neuron populations (e.g., cortex, thalamus), with one caveat. Due to the spread of putamen AAV injection into the globus pallidus (Fig. 2a), there was robust EGFP labeling of neurons in the subthalamic nucleus (STN) across tissue sections for both the native and antibody-amplified reporter (Figs. 2e-f; Extended Data Fig. 3b-c). Except for the STN, endogenous EGFP fluorescence was essentially undetectable outside of the midbrain. Antibody amplified EGFP expression revealed the same pattern of midbrain DA and STN neuron labeling, but with additional sparse cell labeling in the striatum and globus pallidus encompassing the injection site. To avoid misrepresenting the specificity of retrograde DA-neuron labeling in the midbrain, we quantified cells using ROIs that both included and excluded the STN to compare specificity (Fig. 2f). In the original ROIs containing the STN, the specificity of labeling was poor for both endogenous and amplified datasets due to the relatively high number of STN neurons labeled. For the endogenous reporter dataset, we quantified 6,840 anti-TH+ cells and 739 EGFP+ cells in total across sections.

However, 363 of those EGFP+ cells were TH-negative cells detected in the STN (50.88% labeling specificity). In the antibody-amplified reporter dataset, we quantified 12,387 anti-TH+ cells and 2,896 anti-GFP+ cells in total, but again, 1,084 anti-GFP+TH-cells were detected in the STN (62.57% labeling specificity). After excluding cell counts in the STN for each ROI, there remained only 15 EGFP+TH-cells for the endogenous reporter (96.14% labeling specificity) and 147 anti-GFP+TH-cells for the amplified reporter (92.48% labeling specificity). Although the overall specificity of both endogenous EGFP and amplified anti-GFP labeling was low (50.88% and 62.57%, respectively) due to off-target STN transduction, labeling within the midbrain retained high specificity (96.14% and 92.48%) for DA neurons regardless of signal amplification (Fig. 2f). Notably, a local injection of cjDAE8-dTom directly in the anterior midbrain, most proximal to the STN, did not cause off-target labeling in this region (Figs. 2a-b, 3d), likely due to the strong insulating effect of high-density fiber tracts in NHPs. Because of the dose-dependent manner of enhancer-AAV labeling, we believe that off-target STN labeling will only be an issue for mistargeted high-titer local injections or high-titer systemic injections of cjDAE8 AAVs. Nevertheless, this proof-of-concept experiment demonstrates that AAV2retro-cjDAE8 vectors enable robust targeting of projection-defined DA neuron subpopulations in marmoset. Furthermore, intersectional viral strategies can be used to avoid STN labeling altogether, such as AAV2-retro-cjDAE8 driving Cre or tTA expression, combined with midbrain local injection of AAVs driving Cre- or tTA-dependent transgenes.

**Figure 3.**
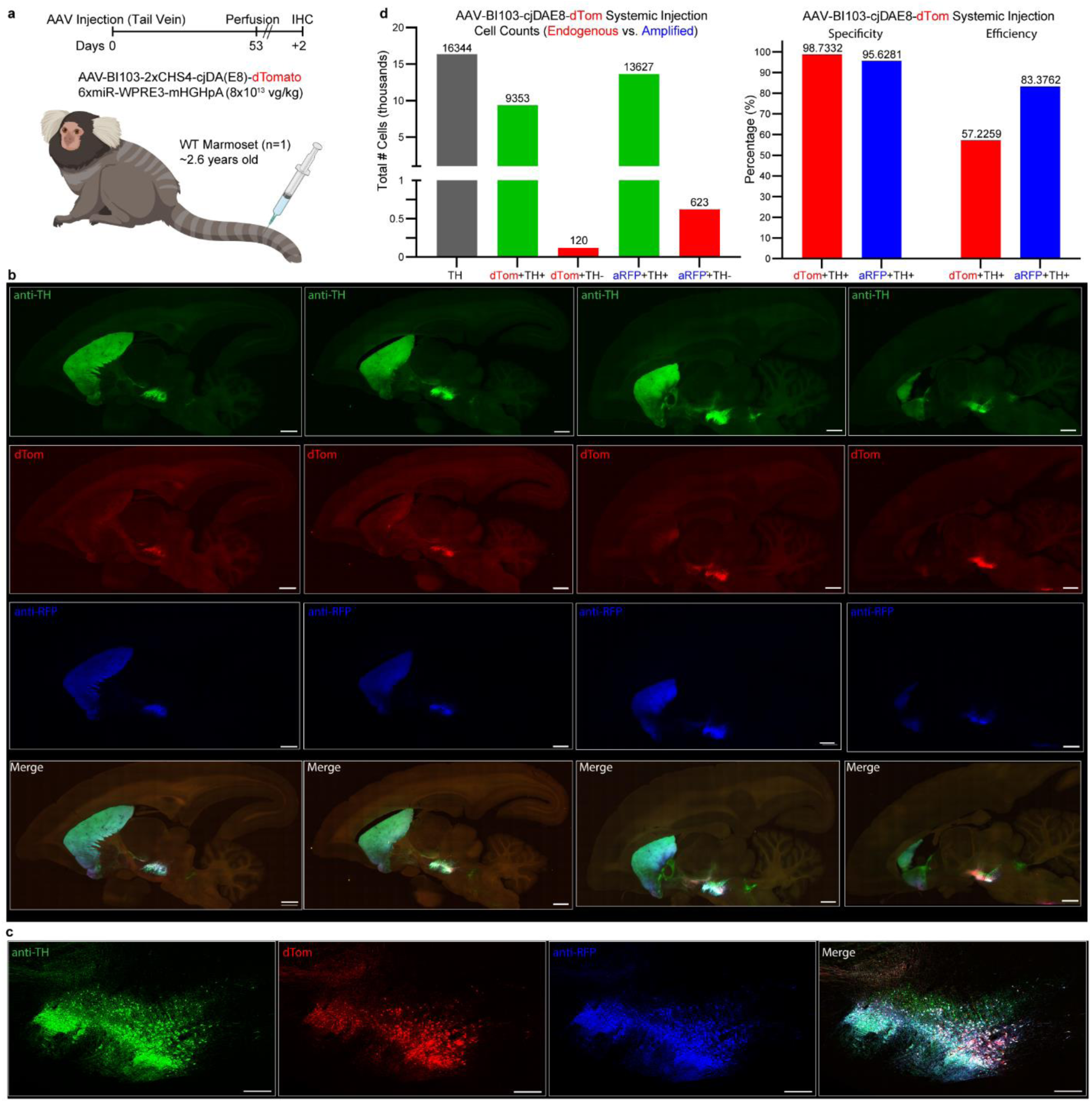
Intravenous cjDAE8-AAV delivery enables selective systemic targeting of marmoset DA neurons. **a,** Timeline and schematic of the systemic injection strategy with AAV-BI103 (teil-vein injeciton, 8×10¹³ vg/kg dose) using a cjDAE8-dTom cassette engineered to de-target peripheral tissues. Schematics made in BioRender. **b,** Representative sagittal sections of the whole-midbrain quantification across serial ROIs. Scale bar, 2 mm. **c,** Higher magnification representative confocal image demonstrating high degrees of colocalization between channels. Scale bar, 500 μm. **d,** (left panel) Aggregated quantification for 14 serial midbrain ROIs showing TH+ neurons and reporter-positive subsets for endogenous (dTom) and antibody-amplified (anti-RFP) conditions: 16,344 TH+; 9,353 dTom⁺TH⁺+; 120 dTom⁺TH⁻; 13,627 anti-RFP⁺TH⁺; 623 anti-RFP⁺TH⁻ cells. (right panel) Summary plots show 98.73% specific and 57.23% efficient DA labeling for endogenous dTom. Similarly, aRFP labeling was 95.63% specific but featured 83.38% efficiency. ***Definitions*.** Specificity = (reporter+TH+ / total reporter+)×100; Efficiency = (reporter+TH+ / total TH+)×100.

### Intravenous cjDAE8-AAV delivery enables selective systemic targeting of marmoset DA neurons

Given the expense, time cost, and technical difficulty of MRI-targeted stereotactic AAV injections, the versatility of targeting DA neurons via local or systemic delivery in NHPs is highly valuable. Considering this and the remarkable specificity of labeling after high-titer local injection, we eagerly tested whether systemic targeting of DA neurons in the marmoset by intravenous AAV injection could achieve robust transgene expression with brain-wide specificity. However, due to the exceedingly high AAV doses required for sufficient CNS expression, dorsal root ganglia toxicity and hepatotoxicity are widely reported complications in primates^58,59^. We therefore modified the 2xCHS4-cjDAE8-dTom construct, replacing the WPRE with 6x total miR binding sites for de-targeting the dorsal root ganglia (DRG, 3x) and liver hepatocytes (3x)^60,61^, followed by the WPRE3 sequence. To enable delivery across the NHP blood-brain barrier (BBB), we packaged the modified construct into AAV-BI103, a capsid variant previously described^49,62^ with efficient BBB penetration in marmosets (Fig. 3a).

High-dose systemic delivery of AAV-BI103-cjDAE8-dTom by tail-vein injection (8E+13 vg/kg) resulted in highly efficient and specific labeling in the midbrain for both native and amplified reporter fluorescence (Fig. 3a-c) with no detectable expression in the DRG and liver (Extended Data Fig. 4). To comprehensively assess the specificity and efficiency of both endogenous and antibody-amplified reporter labeling in midbrain DA neurons, we sampled the entire hemisphere using serial sagittal sections from wells separated by 100 μm. Using generous midbrain ROIs, we quantified 16,344 anti-TH+ cells, among which 9,353 were dTom+TH+ cells (57.23% labeling efficiency, enriched in ventral populations). Remarkably, only 120 dTom+TH-were detected, resulting in 98.73% specific DA labeling (Fig. 3d). As we will discuss later for mouse systemic delivery, endogenous expression at lower systemic doses of cjDAE8 AAVs preferentially labels ventral DA neuron populations, as observed here. Therefore, at higher systemic doses, or more feasibly with a different AAV capsid that exhibits higher BBB transport efficiency, both ventral and dorsal populations are labeled.

After antibody amplification, we detected 13,627 anti-RFP+TH+ cells (83.38% labeling efficiency), a 26% increase over endogenous dTom labeling. Remarkably, only 623 anti-RFP+TH-cells were detected, retaining 95.63% specific DA labeling (Fig. 3d). After antibody amplification, very dim off-target anti-RFP labeling in the STN and relatively strong but uniquely patterned labeling in medial regions superior to the VTA were observed in relevant ROIs and more extensively observed in the 3D-cleared hemisphere. Terminal projections from the off-target cell bodies in these regions were only discernable in the ventral medulla (dTom+ and anti-RFP+, but TH-). Finally, we performed 3D whole-tissue clearing, combined with immunofluorescent staining for reporter and TH expression, to provide the community with an atlas of cjDAE8-driven expression throughout the brain (LifeCanvas Technologies, Supplementary Videos 1-2). These data are also publicly available via NeuroGlancer. Note that, due to antibody costs for achieving robust penetration throughout the whole hemisphere, there was a notable gradient of decreasing fluorescent intensity for anti-RFP and anti-TH labeling observed moving deeper into the tissue from medial to lateral. Consequentially, the most dorsolateral DA-containing ventral midbrain regions were poorly labeled. We therefore performed another round of immunostaining with Fab fragments, which successfully labeled these regions, but further enhanced the decreasing intensity gradient moving deeper into the tissue. This gradient is apparent in the supplementary videos, as we selected constant intensity thresholds throughout the tissue. We therefore recommend interactively viewing this data through Neuroglancer, allowing users to manually adjust the intensity as they move deeper in the tissue. Stacked confocal images of immuno-stained serial sections using the other hemisphere are available on Neuroglancer as well to cross-reference poorly labeled deep regions after immunostaining the cleared hemisphere.

### Systemic and local delivery of cjDAE-AAVs in mice

Next, we characterized the effective dose ranges of cjDAE8-AAV systemic delivery for targeting of DA neurons in mice. Male and female mice received retro-orbital AAV-PHP.eB-2xCHS4-cjDAE8-dTom injections (80 μl) at a dose of either 5E+10 vg (n=4) or 5E+11 vg (n=4) (Fig. 4a). Both AAV doses resulted in robust transduction of midbrain DA neurons that varied in strength and specificity of expression, with the high dose exhibiting weak off-target expression after antibody amplification (Fig. 4b). A two-way repeated-measures ANOVA with factors Dose (5E+10 vg, 5E+11 vg) and Marker (TH+, dTom+TH+, dTom+TH−, anti-RFP+TH+, anti-RFP+TH− cells) revealed a significant Dose×Marker interaction (F(4,24)=8.760, P=0.0002), with main effects of Dose (F(1,6)=10.30, P=0.0184) and Marker (F(4,24)=101.3, P<0.0001).

**Figure 4.**
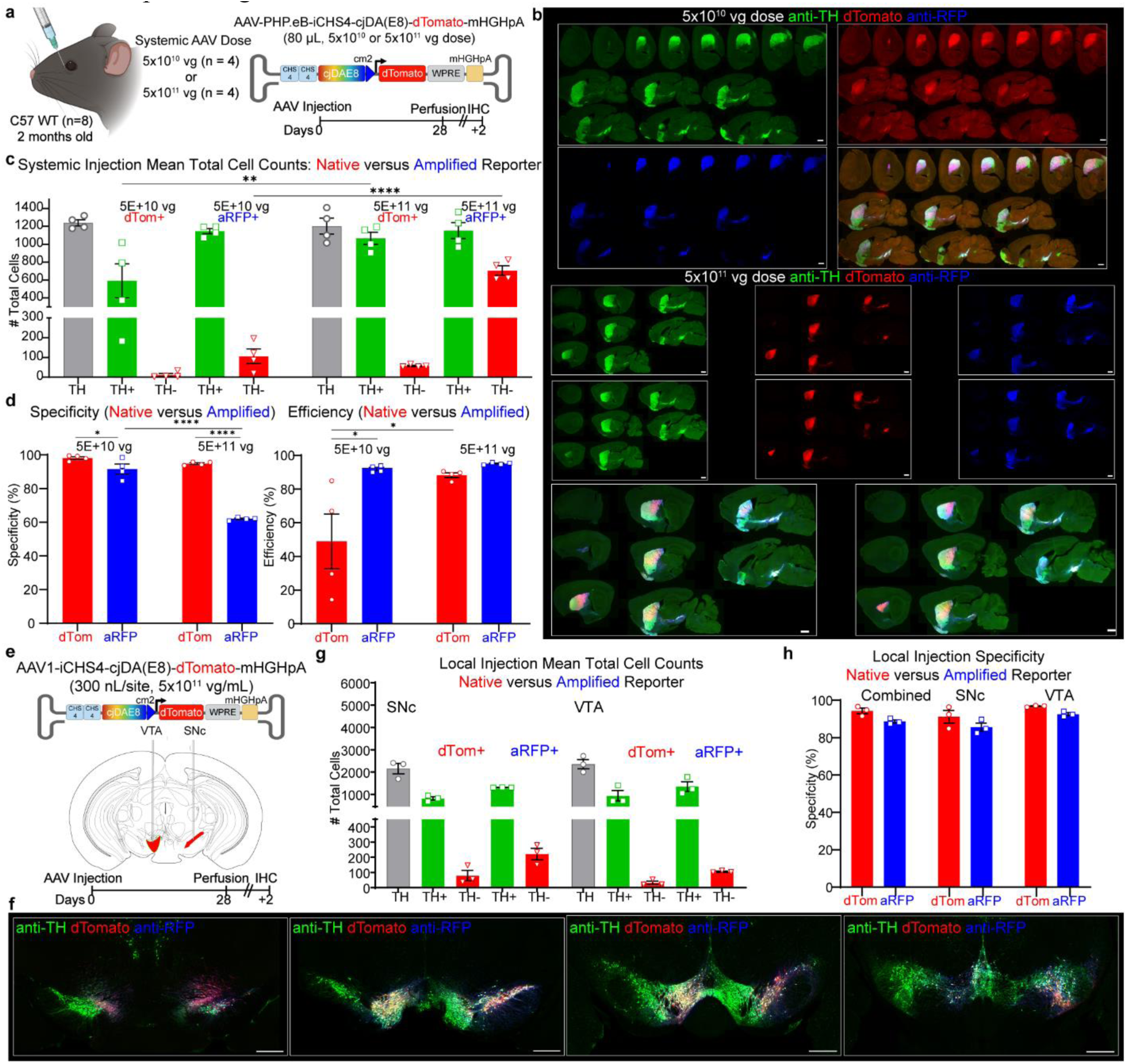
Dose-dependent systemic and local cjDAE8-AAV delivery enables selective targeting of mouse DA neurons. **a,** Male and female C57 WT mice received retro-orbital AAV-PHP.eB–cjDAE8-dTom at 5E+10 or 5E+11 vg (n=4/group) and perfused after four weeks of viral incubation. **b,** Both doses robustly label midbrain DA neurons. At 5E+10 vg, endogenous and amplified signals are highly specific; at 5E+11 vg, endogenous remains specific while amplification reveals sparse or dim off-target labeling. Scale bar, 2 mm. **c,** Two-way RM-ANOVA (Dose×Marker) was significant (F(4,24)=8.760, P=0.0002). High dose increased dTom⁺TH⁺ (+475, adj P<0.001) and anti-RFP⁺TH⁻ (+600, adj P<0.0001); TH⁺, dTom⁺TH⁻, anti-RFP⁺TH⁺ were unchanged (all adj P>0.3). **d,** Specificity showed a strong Dose×Marker interaction (F(1,6)=94.34, P<0.0001). At 5E+10 vg: dTom 98.1% vs anti-RFP 91.6% (adj P=0.0289). At 5E+11 vg: dTom 94.8% vs anti-RFP 62.0% (adj P<0.0001). Across doses, dTom specificity was stable (Δ=−3.29%, P=0.293) while anti-RFP dropped (Δ=−29.6%, P<0.0001). Efficiency showed no interaction (F(1,6)=5.142, P=0.064) but increased with dose (F(1,6)=6.394, P=0.0448) and differed by reporter (F(1,6)=9.744, P=0.0205): at 5E+10 vg, anti-RFP>dTom (92.5% vs 49.0%, adj P=0.0176); at 5E+11 vg, they converged (95.8% vs 88.9%, adj P=0.813). Dose raised dTom efficiency (+39.9%, P=0.0106) but not anti-RFP (Δ=−3.25%, P=0.955). **e,** Male and female C57 WT mice (n=3) were locally injected with AAV1-cjDAE8-dTom (300 nL/site; 5E+11 vg/ml) in the SNc and VTA (n=3), then perfused four weeks later.**f,** Representative confocal images showing robust, specific VTA and SNc labeling along the A–P axis for both endogenous and amplified reporter expression. Scale bar, 500 μm. **g,** As expected, there was a significant Region×Marker interaction (F(8,24)=14.94, P<0.0001): Combined ROI > SNc/VTA for TH⁺, dTom⁺TH⁺, anti-RFP⁺TH⁺. Off-target labeling(dTom⁺TH⁻, anti-RFP⁺TH⁻) did not differ across regions (all adj-P>0.9). **h,** No Region×Reporter interaction, but dTom showed higher overall specificity than anti-RFP (F(1,6)=17.69, P=0.0056), ∼+5.1% on average across Combined/SNc/VTA. Data are mean ± s.e.m.; points are individual mice. ***Definitions.*** Specificity = (reporter⁺TH⁺ / reporter⁺ total)×100; Efficiency = (reporter⁺TH⁺ / TH⁺ total)×100. Asterisks denote significance: * p<0.05, ** p<0.01, *** p<0.001, **** p<0.0001; not significant comparisons were not plotted (two-sided tests; multiple-comparisons corrections as indicated). Mouse and AAV schematics made in BioRender.

Bonferroni post-hoc testing confirmed that, compared to 5E+10 vg, the 5E+11 vg dose increased mean counts for dTom+TH+ (+475 cells, adjusted P<0.001) and anti-RFP+TH− numbers (+600 cells, adjusted P<0.0001), whereas TH+ (Δ=+37 cells), dTom+TH− (Δ=+46), and anti-RFP+TH+ (Δ=+6) cells did not change (all adjusted P>0.3; Fig. 4c). The Dose×Marker interaction was strong (F(1,6)=94.34, P<0.0001) for the calculated specificity of labeling (Fig. 4d), with main effects of Dose (F(1,6)=91.41, P<0.0001) and Marker (F(1,6)=210.7, P<0.0001). At low doses, both native and amplified reporter expression attained high specificity, but with lower labeling efficiency and preferential labeling of ventral DA populations without reporter amplification. After amplification, both dorsal and ventral populations were labeled, with only barely detectable STN labeling in some cases. In contrast, higher doses resulted in both specific and efficient native labeling but significantly reduced specificity after antibody amplification.

Notable regions with detectable dim off-target reporter expression include the STN, lateral mammillary nucleus, and magnocellular nucleus. This finding highlights the dose-dependence of enhancer-AAV performance, with higher doses increasing both expression level in DA neurons and low-level leaky expression.

At 5E+10 vg, there was no significant difference in specificity between endogenous fluorescence and amplified reporter expression (dTom 98.1% vs anti-RFP 91.6%; adjusted P=0.0289). In contrast, at 5E+11 vg, dTom specificity remained high (94.8%), whereas anti-RFP specificity dropped to 62.0% (adjusted P<0.0001). Across doses, specificity remained unchanged for dTom (Δ=−3.29%, adjusted P=0.293), but decreased markedly for anti-RFP (Δ=−29.6%, adjusted P<0.0001). Regarding labeling efficiency (Fig. 4d), there was no Dose×Marker interaction (F(1,6)=5.142, P=0.064), but main effects of Dose (F(1,6)=6.394, P=0.0448) and Marker (F(1,6)=9.744, P=0.0205) were significant. At 5E+10 vg, anti-RFP labeled a greater fraction of TH+ neurons than dTom (92.5% vs 49.0%; adjusted P=0.0176). However, efficiencies converged between dTom and anti-RFP at the 5E+11 vg dose (95.8% vs 88.9%; adjusted P=0.813). Furthermore, as the dose increased, dTom efficiency increased (+39.9%, adjusted P=0.0106), whereas anti-RFP efficiency remained unchanged (Δ=−3.25%, adjusted P=0.955). Together, these results suggest that cjDAE8-AAV is a robust tool for systemic targeting of DA neurons in mice.

Next, we examined the efficacy and specificity of local injection of cjDAE8-AAV in labeling midbrain DA neurons. Male and female mice (n=3 each) received a unilateral injection of AAV1-2xCHS4-cjDAE-dTom (300 nl, 5E+11 vg/ml per site) in the SNc and VTA (Fig. 4e). At this dose, robust and specific reporter labeling was observed throughout the anterior-posterior axis for both endogenous and amplified signals (Fig. 4f). For total cell counts, a two-way repeated-measures ANOVA with factors Region (Combined, SNc, VTA) and Marker (TH+, dTom+TH+, dTom+TH−, anti-RFP+TH+, anti-RFP+TH−) revealed a significant Region×Marker interaction (F(8,24)=14.94, P<0.0001) and main effects of Region (F(2,6)=20.19, P=0.0022) and Marker (F(4,24)=223.2, P<0.0001). As expected, Šídák post-hoc tests showed higher counts in the combined ROI than in SNc and VTA alone for TH+ (Δ=+2457 and +2257 cells; both adjusted P<0.0001), for dTom+TH+ (Δ=+928 and +808 cells; adjusted P=0.0121 and 0.0436), and for anti-RFP+TH+ cells (Δ=+1348 and +1302 cells; adjusted P=0.0001 and 0.0002). dTom+TH− and anti-RFP+TH− did not differ across regions (all adjusted P>0.9; Fig. 4g). With factors Region (Combined, SNc, VTA) and Reporter (dTom vs anti-RFP, both TH+), there was no interaction (F(2,6)=0.1375, P=0.874) and no main effect of Region (F(2,6)=4.533, P=0.0632), but specificity was higher for dTom overall (main effect of Reporter: F(1,6)=17.69, P=0.0056). Across regions, dTom specificities were 94.3% (Combined), 91.2% (SNc), and 96.8% (VTA), compared to 88.7%, 85.7%, and 92.6%, respectively, for anti-RFP (Fig. 4h); the average dTom–anti-RFP difference was +5.14% (SE 1.22%; 95% CI 2.15–8.14). These results indicate that local delivery of cjDAE-AAV at this dose enables efficient and specific targeting of mouse SNc and VTA DA neurons, even after antibody-amplification across regions.

### cjDAE8-AAV-GCaMP6f enables imaging of SNc DA terminals in the striatum

To demonstrate the utility of cjDAE8-AAV for *in vivo* calcium recordings of DA neurons, we bilaterally injected AAV9-cjDAE8-GCaMP6f in the SNc of mice (n=4) (8E+12 vg/ml). Optic fiber cannulas were implanted into either the central striatum (CS) or dorsolateral striatum (DLS) of each hemisphere to compare CS and DLS DA activity simultaneously with fiber photometry (Fig. 5a, left panel). Post-hoc histology confirmed specific expression of GCaMP6f in DA neurons (Fig. 5a, right panel). CS and DLS DA axons were targeted due to the distinct functional specialization of DA terminal activity in these subregions reported previously: the DLS (movement) and the CS (reward, punishment, salience)^45,54,55^. While mice were head-fixed to a self-driven treadmill, photometry recordings were acquired i) over two sessions with synchronized measurements of instantaneous velocity to capture spontaneous locomotion bouts and ii) over two sessions of 9x trials delivering an unconditioned and salient air-puff stimulus.

**Figure 5.**
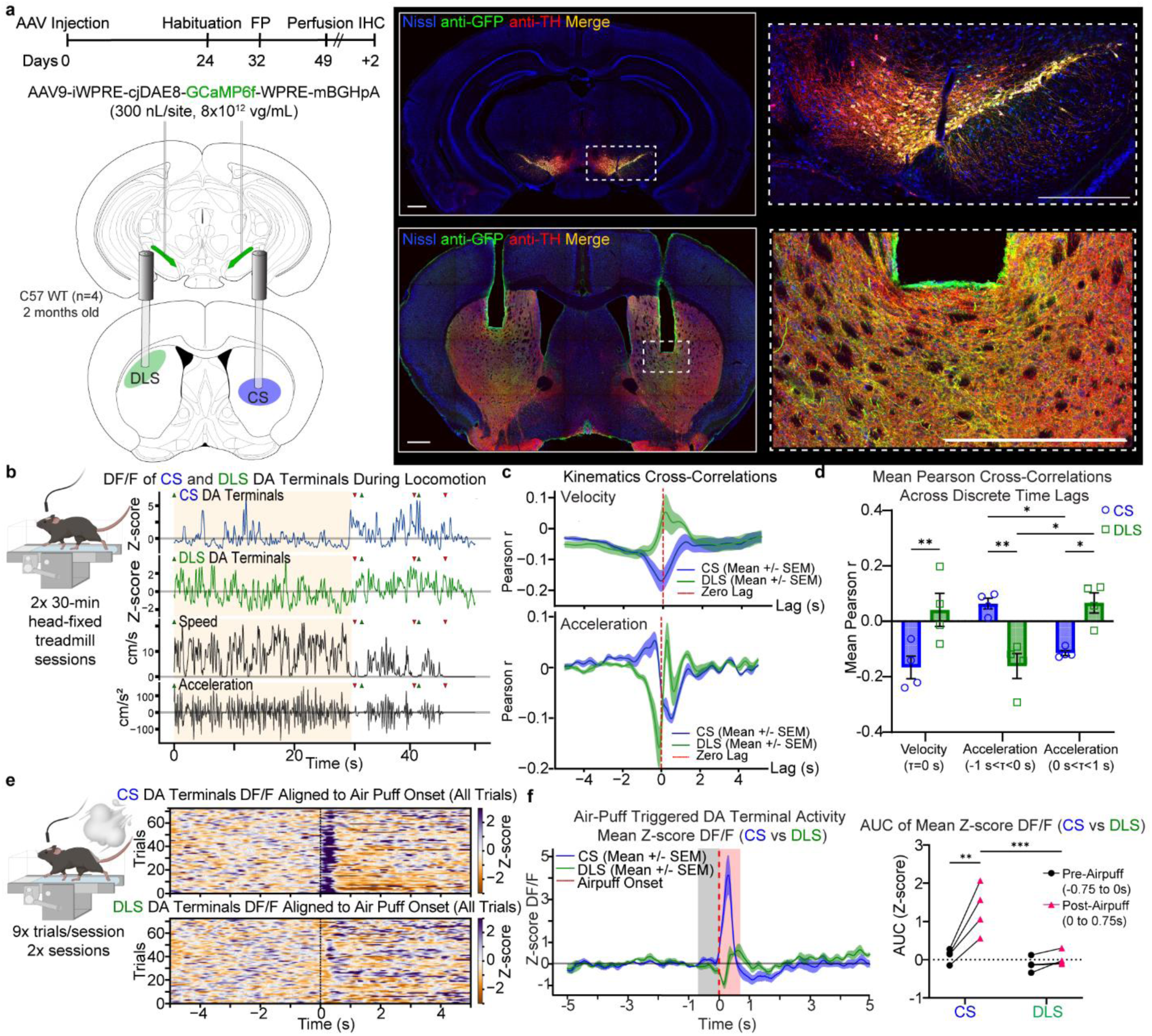
SNc DA terminals in CS and DLS differentially encode movement kinematics and salient stimuli. **a,** C57 WT mice (n=4) received bilateral SNc injections with AAV9–cjDAE8–GCaMP6f (8×10¹² vg mL⁻¹) and implanted with fibers in the CS and DLS. Scale bar, 500 μm. **b,** Z-scored CS/DLS signals were examined during spontaneous movement bouts (speed >5 cm s⁻¹; |accel|>50 cm s⁻²), showing distinct activity patterns. **c,** CS and DLS DA terminals oppositely correlate with velocity and acceleration. Velocity at τ=0 (top): DLS positive (r=+0.0412 ± 0.0600), CS negative (r=−0.1663 ± 0.0404). Acceleration mirrors at negative versus positive time lags (bottom): CS peak at −0.50 ± 0.14 s (r=+0.0635 ± 0.0194) and trough at +0.44 ± 0.08 s (r=−0.1144 ± 0.0090); DLS trough at −0.11 ± 0.02 s (r=−0.1614 ± 0.0450) and peak at +0.47 ± 0.18 s (r=+0.0662 ± 0.0365). **d,** Summary of mean Pearson r across time lags for velocity and acceleration. RM-ANOVA Region×Period interaction (F(2,6)=23.73, P=0.0014). Šídák: τ=0, CS<DLS (Δ=−0.208, adj P=0.0059); Accel −1<τ<0, CS>DLS (Δ=+0.225, adj P=0.0040); Accel 0<τ<+1, CS<DLS (Δ=−0.181, adj P=0.0109). Within-region: CS Accel(−)>Accel(+) (Δ=+0.178, P=0.0346); DLS Accel(+)>Accel(−) (Δ=+0.228, P=0.0113). **e,** Onset-aligned mean Z-score fluorescence changes show rapid CS activation across all air puff trials, with minimal response in the DLS. **f,** AUC analysis of mean Z-score fluorescence using pre- and post-onset windows (±0.75 s). RM-ANOVA Region×Time (F(1,6)=17.82, P=0.0056). CS increases pre→post (0.109 ± 0.091 → 1.312 ± 0.324; adj P=0.0010); DLS no change (−0.122 ± 0.094 → 0.019 ± 0.096; adj P=0.9174). Post period: CS>DLS (mean diff = 1.293; adj P=0.0006). Data are mean ± s.e.m.; points are individual mice. Asterisks denote significance: * p<0.05, ** p<0.01, *** p<0.001, **** p<0.0001; not significant comparisons were not plotted (two-sided tests; multiple-comparisons corrections as indicated). Mouse-related schematics made in BioRender.

During periods of spontaneous movement, calcium transients of DA CS and DLS terminals showed distinct activity patterns (Fig. 5b). Pearson cross-correlations of Z-score fluorescence with velocity and acceleration during movement periods (>5 cm/s, >50 cm/s^2^) revealed that CS and DLS DA-terminal activity are inversely correlated with each other during spontaneous locomotion. At zero-time lags (τ=0) for cross-correlations with velocity, mean Pearson r values (mean ± s.e.m.) were positive for DLS DA fluorescence (0.0412 ± 0.0600), but negative for CS DA fluorescence (−0.1663 ± 0.0404). In other words, higher and lower speeds are associated with greater activity in DLS and CS DA terminals, respectively (Fig. 5c). Similarly, Pearson cross-correlations with acceleration across time lags for the two subregions mirrored each other. With symmetry across zero-time lags, Pearson r values at negative time lags (mean ± s.e.m.)

Contained a deep trough for the DLS (−0.1614 ± 0.0450; lag −0.110 ± 0.019 s) and a peak for the CS (0.0635 ± 0.0194; lag −0.500 ± 0.144 s). In contrast, there was a trough for the CS (−0.1144 ± 0.0090; lag 0.440 ± 0.077 s) and peak for the DLS (0.0662 ± 0.0365; lag 0.470 ± 0.178 s) at positive lags (Fig. 5c). The trough at τ<0 and peak at τ>0 for DA terminals in the DLS suggests that activity increases before deceleration and after acceleration start, respectively. In contrast, the trough at τ>0 and peak at τ<0 for DA terminals in the CS suggest that activity increases before acceleration and after deceleration.

We further quantified mean cross-correlations between DA-terminal signals in the CS and DLS and kinematics (velocity, acceleration) across mice (Fig. 5d). A two-way repeated-measures ANOVA with factors Region (CS, DLS) and Time-Lag Period (Velocity τ=0; Acceleration −1<τ<0 or Acceleration 0<τ<+1) revealed a significant Region×Period interaction (F(2,6)=23.73, P=0.0014) and a main effect of Region (F(1,3)=14.41, P=0.0321), with no main effect of Period (P=0.7477). Šídák-corrected post-hoc analysis showed that: at velocity τ=0 s, CS correlations were more negative than DLS (mean CS −0.166, DLS +0.041; Δ=−0.208, 95% CI −0.329 to −0.086, adjusted P=0.0059). For cross-correlations with acceleration at negative time lags (−1<τ<0 s), CS mean Pearson r was more positive than DLS (CS +0.064, DLS −0.161; Δ=+0.225, 95% CI +0.103 to +0.347, adjusted P=0.0040). In contrast, at negative time legs (0<τ<+1 s), CS mean Pearson r was more negative than DLS (CS −0.114, DLS +0.066; Δ=−0.181, 95% CI −0.302 to −0.059, adjusted P=0.0109). Within-region comparisons mirrored this sign flip: for CS, Accel(−) > Accel(+) (Δ=+0.178, P=0.0346), and for DLS, Accel(+) > Accel(−) (Δ=+0.228, P=0.0113). In summary, at zero lag, CS DA activity negatively correlated with velocity, while DLS positively correlated with velocity. Furthermore, CS DA activity positively correlated with acceleration at negative time lags (−1<τ<0) and became negatively correlated at positive time lags (0<τ<+1), whereas DLS showed the opposite polarity. Together with the significant interaction, these data further corroborate the known region-specific, time-locked coupling of DA-terminal activity to movement kinematics^55,63^.

DA terminals in the CS and DLS also differentially responded to salient air-puff stimuli. Triggered mean Z-score fluorescence plots aligned to air-puff onset (t=0) across trials robustly confirmed opposing roles of CS and DLS DA terminals in the encoding of salient stimuli (Fig. 5e). To quantify activity changes after stimulus onset, area under the curve (AUC) analysis was performed on the mean Z-score trace for each mouse using a 0.75-s window before and after stimulus onset (Fig. 5f). Two-way repeated-measures ANOVA with factors Region (CS, DLS) and Time (Pre, Post) revealed a Region×Time interaction (F(1,6)=17.82, P=0.0056), with significant main effects of Region (F(1,6)=11.64, P=0.0143) and Time (F(1,6)=28.53, P=0.0018). CS DA terminals showed robust air-puff-evoked activity (mean AUC ± s.e.m.) from pre (0.109 ± 0.091) to post (1.312 ± 0.324) air-puff delivery (Bonferroni t(6)=6.762, adjusted P=0.0010), whereas DLS terminals did not (–0.122 ± 0.094 vs 0.019 ± 0.096; t(6)=0.792, adjusted P=0.9174), yielding a significant Region×Time interaction (F(1,6)=17.82, P=0.0056) and a marked CS>DLS activity difference (mean diff = 1.293, 95% CI [0.637, 1.949], t(12)=5.044, adjusted P=0.0006) in the post-air puff period. Together, these data further corroborate prior evidence that DA terminals in the CS (Vglut2+ and Calb1+) but not in the DLS (ANXA1+) are rapidly activated in response to salient stimuli^55^.

### AAV-cjDAE8-channelrhodopsin enables optogenetic self-stimulation of VTA DA neurons in an operant behavioral paradigm

Finally, to demonstrate the utility of cjDAE8-AAV for in vivo functional manipulation of DA neurons, we bilaterally injected in the VTA (300 nl per site) of male (n = 4) and female (n = 6) mice with AAV1-cjDAE8-ChR2(H134R)-EYFP (n = 6) or AAV1-cjDAE8-dTom (n = 4; 1E+12 vg/ml) (Fig. 6a–b). To span an effective dose range for ChR2, we tested 1E+13 (n = 1), 5E+12 (n = 1), 2E+12 (n = 2), and 1E+12 vg/ml (n = 2). After ≥14 days of expression, naïve mice were trained for three sessions in a self-paced VTA-DA optogenetic self-stimulation task (active vs inactive nose-poke; 2 s, 20 Hz, 4.5–5 mW opto-stim per active poke; max 180 stimulations per 30 min; Fig. 6c).

**Figure 6.**
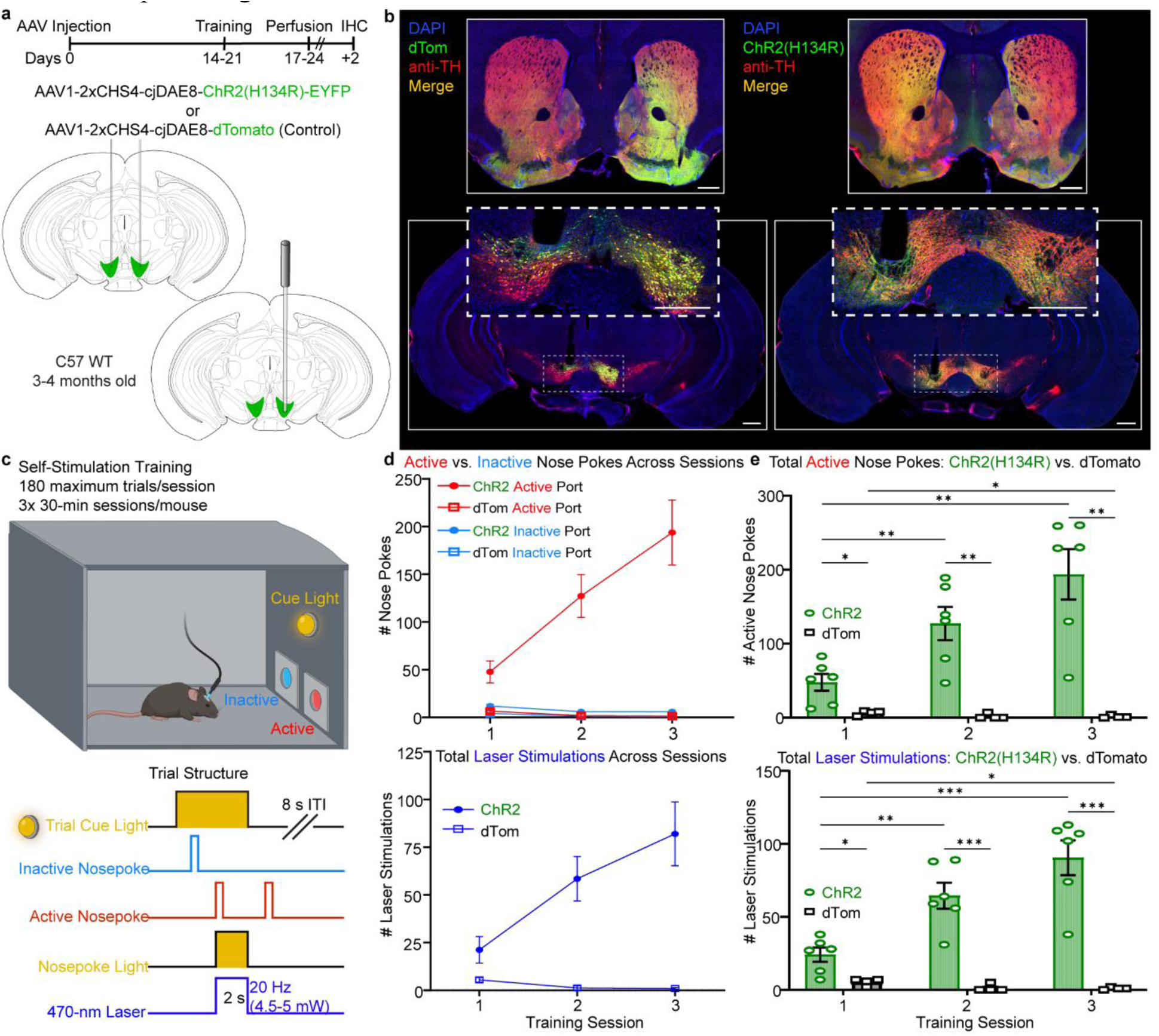
Optogenetic stimulation of VTA DA neurons elicits robust self-stimulation behavior in mice. **a,** Bilateral VTA injections (300 nl/site) of AAV1-cjDAE8 driving ChR2(H134R)-EYFP (n=6, 1E+12 to 1E+13 vg/ml) or dTomato (n=4, 1E+12 vg/ml). **b,** Representative expression of either construct at the injection site. **c,** Optogenetic self-stimulation task: cue-light, active vs inactive ports; active poke triggers 2-s, 20 Hz laser (4.5–5 mW) and 8-s ITI (max 180 stim/session); mouse-related schematic made in BioRender. **d,** Learning curves across three sessions: active/inactive nose-pokes (top) and delivered stimulations (bottom; same measure summarized in **e**). Three-way RM-ANOVA (Session × Transgene × Port) showed significant main effects and interactions (all P≤0.0023); compared to inactive, active nose pokes escalated across days in ChR2 mice (all Šídák-adjusted P ≤0.002) but not in controls (all Šídák-adjusted P≥0.05) **e,** Summary of active nose-pokes (top) and stimulations (bottom) across training sessions. Two separate two-way RM-ANOVAs (Session × Group) showed Session×Group interactions for active pokes and stimulations (P=0.0006 and 0.0003, respectively) with ChR2 > dTom at each session (Tukey, all P≤0.015). Data are mean ± s.e.m.; points are individual mice. Asterisks denote significance: * p<0.05, ** p<0.01, *** p<0.001, **** p<0.0001; not significant comparisons were not plotted (two-sided tests; multiple-comparisons corrections as indicated).

Across training (Fig. 6d), a three-way ANOVA (Session × Transgene × Port) on nose-poke counts revealed significant main effects of Session (F(2,16)=13.66, P=0.0003), Transgene (ChR2 vs dTom; F(1,8)=20.63, P=0.0019) and Port (Active vs Inactive; F(1,8)=20.23, P=0.0020), as well as Session×Transgene (F(2,16)=17.22, P=0.0001), Session×Port (F(2,16)=15.99, P=0.0002), Transgene×Port (F(1,8)=19.42, P=0.0023), and a three-way interaction (F(2,16)=16.88, P=0.0001). Simple-effects tests showed that in ChR2-expressing mice, nose-poking of the active port vigorously increased across sessions (Sídák-adjusted: Day1 vs Day2 P=0.0003; Day1 vs Day3 P<0.0001; Day2 vs Day3 P=0.0020), whereas nose-poking of the inactive port remained low. In striking contrast, dTom controls responded minimally to either port across sessions (all adjusted P≥0.05).

We further compared group differences in active nose-poke responses and laser stimulations across training (Fig. 6e) with separate two-way RM ANOVAs (Session × Group), using Geisser–Greenhouse correction. For active nose-pokes (Fig. 6e, top), we observed a Session×Group interaction (F(1.472,11.78)=17.15, P=0.0006), main effects of Session (F(1.472,11.78)=14.90, P=0.0011) and Group (F(1,8)=20.06, P=0.0021). Tukey-adjusted post-hoc tests confirmed dramatically higher active-port responses in ChR2-than dTom-expressing mice across all sessions (Day1 P=0.0150; Day2 P=0.0025; Day3 P=0.0024) and escalating responses across days within ChR2 (Day1→Day2 P=0.0058, Day1→Day3 P=0.0052; Day2→Day3 borderline, P=0.0532), with no significant changes across days in controls but a trend of decreasing responses across days (all adjusted P≥0.053). For delivered laser stimulations (Fig. 6e, bottom; note same data from Fig. 6d), we again found a strong Session×Group interaction (F(1.628,13.02)=24.61, P=0.0003) with main effects of Session (F(1.628,13.02)=18.51, P=0.0004) and Group (F(1,8)=34.89, P=0.0021).

Although assessing cellular specificity is more challenging with ChR2, we note that 300 nl injections at titers >2E+12 vg/mL produced visible TH-negative reporter labeling near the injection site in either cell bodies or neurites (see 5E+12 vg/mL for example, Extended Data Fig. 5). Nevertheless, the behavioral phenotype was robust and consistent across animals and titers.

Together, these data indicate that local delivery of cjDAE8-AAV—especially at lower titers—enables selective and robust manipulation of DA circuits *in vivo*.

## Discussion

Through single-nucleus multiomic profiling of marmoset midbrain DA neurons combined with rapid *in vivo* candidate screening across species, we identified a novel gene regulatory element (cjDAE8) that can be used in AAV vectors to potently and specifically drive transgene expression in DA neurons in marmoset and mice. A common obstacle with enhancer-AAV technology is the low-level leaky transgene expression in off-target cell types or brain regions, thereby restricting delivery methods or requiring intersectional viral strategies^37,38,40,41,43,64–67^. We further modified the enhancer-AAV backbone to contain tandem copies of CHS4, a known suppressor of enhancer activity^52^, for insulating the enhancer and minimal promoter from being activated by the upstream ITR sequence^68–70^. To improve expression strength, particularly in NHPs, we replaced mBGHpA with mHGHpA. Using these core elements in the construct backbone, with additional alterations depending on experiment, we systematically validated the performance of cjDAE8-AAVs in mice and marmosets, achieving robust transgene expression with unprecedented specificity, regardless of administration method in both species. All histology data from the mouse and marmoset validation experiments are publicly available and hosted on Neuroglancer for a year. Within the Neuroglancer viewer, users can interact with all high-resolution histology datasets described here for specificity and efficiency analysis after local and systemic delivery. As mentioned previously, we are also hosting the entire dataset for the 3D-cleared hemisphere with immunofluorescent staining, providing a comprehensive overview of expression after systemic delivery.

To further demonstrate the utility of cjDAE8-AAVs, we performed functional validation experiments by locally injecting AAV constructs with cjDAE8 directly driving the expression of transgenes to record (GCaMP6f) or stimulate (ChR2(H134R)) DA neurons in mice. Using fiber photometry, we recorded divergent calcium transients in DLS and CS DA terminals during movement and in response to salient stimuli. In line with prior studies using DAT-Cre or DA-subtype Cre-driver lines, we report that CS and DLS DA-terminal activity were differentially correlated with movement kinematics. Furthermore, DA terminals in the CS, but not DLS, exhibited robust transient activity in response to salient, unconditioned air-puff stimuli^55,63,71,72^.

Finally, using an operant task with responses reinforced by optogenetic stimulation of VTA DA neurons, we elicited robust self-stimulation behavior in ChR2-expressing mice, but not dTomato-expressing controls. These results are highly consistent with those previously reported using a comparable operant optogenetics task in DAT-Cre or TH-Cre rodents^73–76^. Together, these results highlight the power of using cjDAE8-AAVs for functional applications across species. The constructs characterized here and others will become publicly available on Addgene to provide researchers with a comprehensive AAV toolbox supporting applications for DREADDs, optogenetics, monosynaptic rabies tracing, and calcium recordings.

To the best of our knowledge, cjDAE8-AAVs represent the first robust, versatile toolkit for selectively labeling, tracing, recording, and manipulating DA neurons in NHPs with unprecedented specificity and efficiency. In rodents, widely available transgenic lines are the gold standard for selective targeting of DA neurons. Most notably, the DAT-IRES-Cre mouse line achieves greater than 90% specificity of transgene expression by delivering Cre-dependent expression cassettes^22–27,29^. However, the costs of maintaining transgenic rodent colonies (purchasing/breeding, housing, genotyping, etc.) limit access to this powerful tool. In addition, the genetic engineering of these lines to insert Cre (which confers toxicity a high levels) into the genome can disrupt endogenous transcriptional regulation, with unanticipated consequences.

Disrupted gene expression may alter local cellular physiology and circuit-level activity. Indeed, DAT protein expression is reduced in DAT-Cre mice (∼17% in heterozygotes; ∼50% in homozygotes)^28^. Off-target Cre expression in nondopaminergic populations of limbic brain regions (amygdala, septum, hypothalamus, habenula)^29^ and abnormal behaviors (baseline hyperactivity; reduced locomotion/stereotypy after amphetamine)^30^ are observed in DAT-Cre mice as well. These features raise concerns about whether experiments using DAT-Cre (and recombinase-driving transgenics generally) are studying naturalistic cellular and circuit function. Thus, even in rodents, we believe cjDAE8-AAVs offer a cost-effective alternative approach for selectively targeting midbrain DA neurons.

We urge caution when selecting enhancer-AAV doses across experimental purposes and administration routes. After systemic administration in mice, lower doses offer high brain-wide specificity for both native and amplified reporter expression, but unamplified expression may suffer from dim labeling and low efficiency. At the other end, higher doses offer robust, high-efficiency, and specific native transgene expression, but result in low-level leaky expression that is apparent after antibody amplification. Notably, however, the efficiency of antibody-amplified labeling was comparable between doses. For experiments requiring high brain-wide specificity of transgene expression and not high endogenous expression levels (e.g., reporter, recombinase, tTA, TVA), we recommend very low to low systemic doses in mice. Considering that off-target leaky expression is almost always weaker than on-target, for experiments utilizing functional transgenes (e.g., fluorescent calcium sensors, opsins, DREADDs, we recommend medium to high doses to achieve sufficient endogenous expression levels for optimal performance, with some caveats.

In our hands, genetically encoded calcium sensors require high AAV doses for sufficient signal-to-noise ratios due to lower expression levels compared to fluorescent reporters. Furthermore, due to comparatively lower expression levels in off-target labeled cells, contaminating GCaMP signals is not an issue for calcium recording techniques with bulk-signal measurements (e.g., fiber photometry). For experiments using optogenetics and DREADDs, spatially targeted methods involving implantation of optic fiber or infusion cannulas directly over midbrain DA neurons or terminal projection targets largely overcome the potentially confounding effects of low-level off-target expression in distant brain regions. Circumventing AAV transduction in regions susceptible to off-target expression, we demonstrated that direct delivery of cjDAE8-AAVs by local injection enables selective and robust transgene expression across species.

Nevertheless, local injection still requires careful AAV dosing; we observed increasingly lower specificity in mice at titers greater than 2E+12 vg/ml (300-nl injection volume) to express ChR2(H134R)-EYFP (see Extended Data Fig. 5). Considering that enhancer-AAV technology acts through enhancer-minimal promoter interactions that are gated at the DNA level, we also caution against using low-quality AAV preparations with impaired capsid integrity or significant genome truncations. In our experience, this resulted in failed, weak, or even off-target transgene expression in some cases. To ensure accurate dosing and reproducible performance, we therefore stress the importance of using validated, high-quality AAV preparations with low levels of truncated variants, as confirmed by AAV whole-genome sequencing.

## Methods

### Animals

#### Marmosets

Male and female common marmosets (n=4, 2-3 years old) were used for NHP experiments. Marmosets were pair or group-housed in spacious holding rooms maintained on a regular 12-h day/night cycle with controlled temperature (23-28°C) and humidity (40-72%). Perches and enrichment devices were provided in all enclosures. Veterinary staff from the MIT Division of Comparative Medicine (DCM) and researchers provided regular health and behavioral assessments. All animal procedures were approved by the MIT Committee for Animal Care (CAC) and followed veterinary guidelines.

### Mice

Male and female C57BL/6 mice of specified ages were purchased (Jackson Labs, stock # 000664) for rodent experiments. Mice were group-housed with enrichment in a temperature-controlled colony room maintained on a regular 12-h day/night cycle. Food and water were provided *ad libitum*. All animal procedures were approved by the MIT CAC.

### Multiomic analysis and enhancer nomination

#### Marmoset tissue specimen preparation for snRNA and snATAC sequencing

In accordance with MIT CAC protocol, one female marmoset (19-184, aged 2.7 years) was initially sedated and induced with alfaxalone (12 mg/kg) and midazolam (0.3 mg/kg), followed by additional sedation with alfaxalone (4 mg/kg) before euthanasia with a fatal overdose of sodium pentobarbital. The marmoset was then perfused transcardially with ice-cold HEPES buffer containing (in mM): NaCl (110), HEPES (10), glucose (25), sucrose (75), MgCl_2_ (7.5), KCl (2.5) for rapid whole-brain extraction. Regions of the midbrain containing DA neurons were bilaterally micro-dissected from 2-mm tissue slabs. While fresh, the dissected tissue was immediately processed for nuclei isolation and sequencing following standard 10X protocols.

#### 10X snRNA and snATAC-sequencing and analysis

The multome data was generated from 2 reactions using Single Cell Multiome ATAC + Gene Expression v1 chemistry and processed with Cell Ranger ARC 2.0. Unsorted cell nuclei underwent Gel-Bead-in-emulsion (GEM) barcoding and library preparation following the 10X protocol (Extended Data Fig. 1A). A nucleus was included for analysis if it had more than 1,000 genes, between 1,000 and 100,000 chromatin peaks, high transcriptional-start site enrichment (>1), and low nucleosome signal detection (< 2). Out of the 39,767 total cells sequenced, a total of 32,070 passed the quality check. For reaction 1, the median number of ATAC high-quality fragments per cell was 18,482, and the median number of genes expressed per cell was 1,660.

Similarly, for reaction 2, the median number of high-quality fragments per cell was 17,853, and the median number of genes per cell was 1,767. The multiome data from both reactions were combined and annotated with Callithrix_jacchus_cj1700_1.1 (calJac4 in UCSC browser). For joint RNA–ATAC clustering (Fig. 1A), we performed SCTransform normalization on RNA counts and TF–IDF normalization with SVD on ATAC peaks, followed by weighted nearest neighbor (WNN) analysis integrating PCA and LSI reductions using Seurat^77^. Cell-type annotations of the clusters were performed by using canonical marker genes for DA and non-DA populations used in previous studies^78,79^. Reactions 1 and 2 each yielded 141 and 117 DA cells, respectively (n=258). We subclustered additional relevant populations to serve as control populations for enhancer nomination. Specifically, cortical excitatory neurons and inhibitory striatal projection neurons (SPNs) from contaminating dissected tissue were clustered by ADARB2+/KCNA1+ and Meis+ marker expression, respectively. Lymphocytes (B-cells) were clustered by the characteristic PAX5+/TMEM132C-marker. Finally, nigral and VTA inhibitory projection neurons were further subclustered from inhibitory neurons based on the expression of GAD2+/RIT2+/TH-markers.

#### Enhancer candidate nomination

To nominate enhancers, we first generated a consensus of peaks by disjoining overlapped ATAC peaks between the two reactions, then identified differentially accessible chromatin regions (enhancer candidates) enriched in dopamine neurons compared with all other cell types, using logistic regression with sequencing depth as a covariate. Thresholding was set such that only peaks accessible in at least 30% of DA cells and with read counts ≥1.5 log2 fold-change compared to other cell types were considered. The candidate list was further refined by considering genes that are closest to the peaks in linear sequence distance and their mRNA enrichment in DA neurons, together with cross-species sequence conservation and genome context, we selected the top 12 candidates for *in vivo* testing.

### Cloning of enhancer plasmid AAV constructs

For our candidate screening with minimally purified AAV, 12 candidates (DAE1-DAE12) were amplified from purified marmoset genomic DNA (Table 1) and flanked with homologous-overlap sequences for Gibson assembly using enhancer-specific primers and KOD Xtreme DNA Polymerase (Sigma-Aldrich, Cat. # 71975-M). The amplified enhancer candidates were cloned into enhancer-AAV plasmid vectors (eAAV) described previously^43,49^ using restriction enzymes (New England Biolabs) and the NEBuilder HiFi DNA Assembly Cloning Kit (NEB Cat. # E5520S). Between AAV ITR sequences, the expression cassette contained an inverted Woodchuck hepatitis virus Post-transcriptional Regulatory Element (WPRE) as an insulator, the amplified enhancer sequence, a cytomegalovirus (CMV) promoter minimal promoter (CMVmini2)^43^, EGFP or dTomato reporter, WPRE, and a modified bovine growth hormone poly A (mBGHpA). Following the standard transformation protocol, homemade Stbl3 cells (50 μL) were transformed with 2 μL of Gibson assembly product for each cloned enhancer pAAV construct. Mini-preparations of the clones were carried out using the ZR Plasmid Miniprep - Classic Kit (Cat. # D4016). Positive clones were identified by restriction enzyme digestion, followed by whole-plasmid sequencing to confirm 100% alignment to the designed sequence. Confirmed clones were grown in 300-ml LB cultures, and the plasmids were extracted and purified using the NucleoBond Xtra Midi EF Kit (Macherey-Nagel, Cat. # 740420.50). The purified plasmids were sequenced again before AAV production.

**Table 1.**
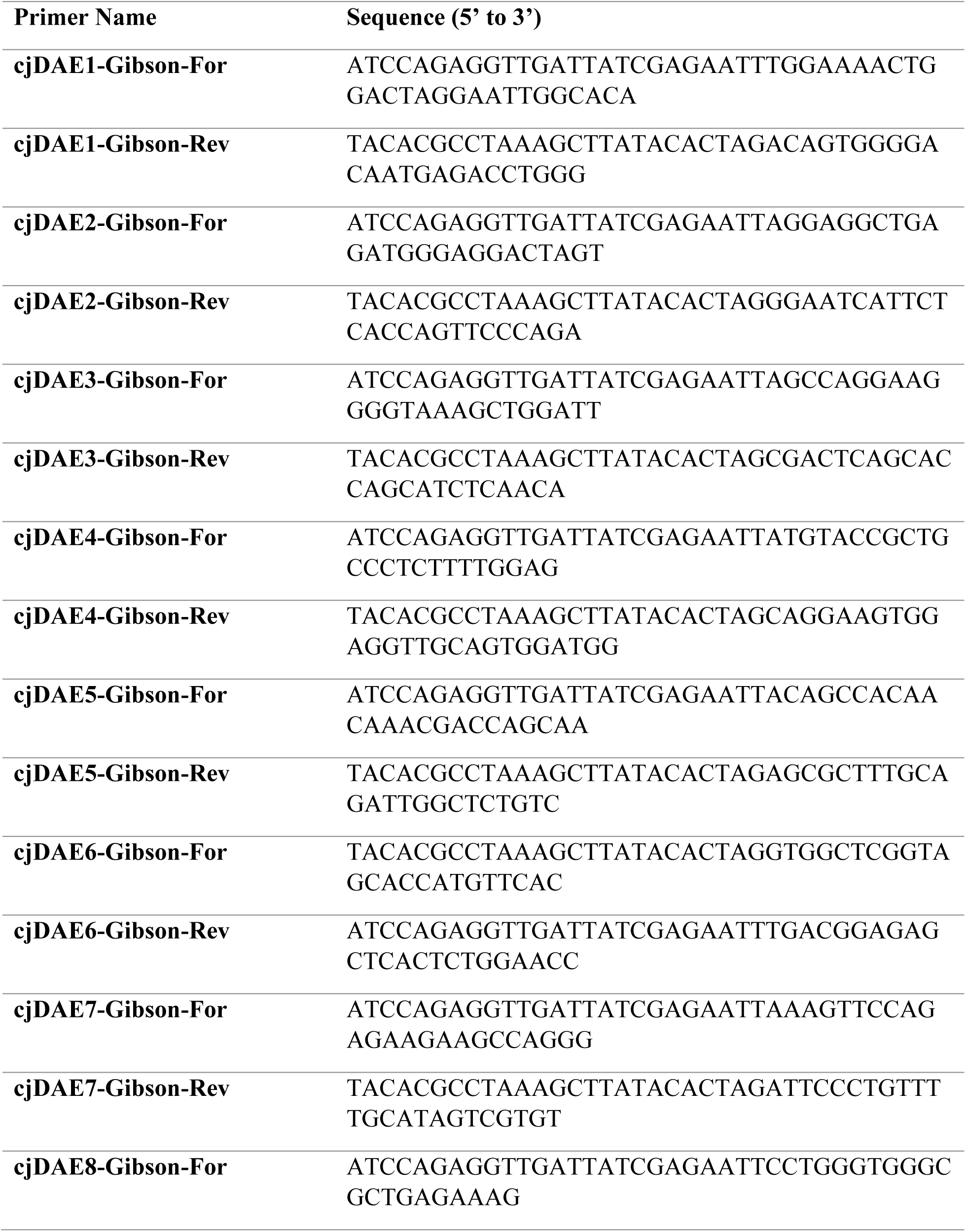
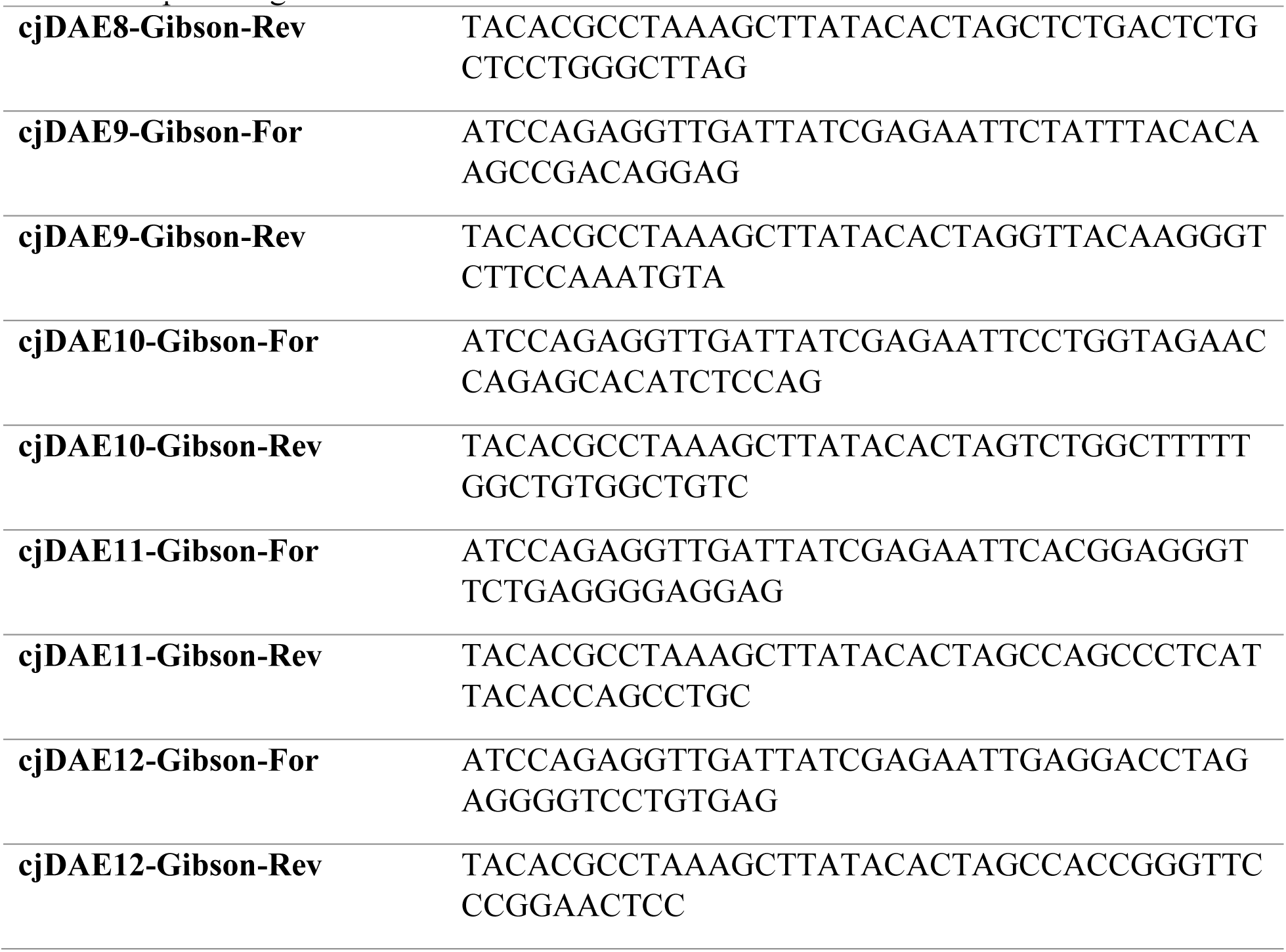
Amplification of candidate DA-enhancer peaks from marmoset gDNA for Gibson assembly of pAAV vectors.

For the best enhancer candidate, cjDAE8, eAAV constructs were further modified to contain a tandem CHS4 insulator^51,52^, different transgenes, downstream WPRE or modified WPRE (WPRE3), and a modified human growth hormone poly A (mHGHpA). For marmoset systemic injection with AAV-BI103, the WPRE was replaced with WPRE3 and 6x miR binding sites to de-target the liver (3x mmu-miR122-5p sites) and dorsal root ganglion (3x hsa-miR-183-5p sites)^60,61^. All eAAV-construct modifications were generated following standard molecular cloning protocols described previously.

### AAV production

Enhancer AAVs for the *in vivo* candidate screen in mice were packaged in-house with AAV2/PHP.eB using a non-titered, minimal purification approach described previously^50^. Preparations of AAV used for local and systemic injection validation, fiber photometry, and optogenetics experiments were packaged either in-house (AAV2/9, AAV2-retro, or AAV-BI103) or at the Boston Children’s Hospital Viral Core (AAV2/1, AAV2/PHP.eB).

#### AAV production with minimal purification

As mentioned previously, we used a shortened purification method^50^ to rapidly screen enhancer candidates. In short, HEK 293 T cells grown in Dulbecco’s Modified Eagle Medium (D-MEM; 11995-040, Gibco, Thermo Fisher Scientific) supplemented with 10% fetal bovine serum were triple-transfected with our AAV enhancer plasmid, pAAV-PHP.eB (Addgene # 103005) and pHelper (Agilent, Cat. # 240071) using Polyethylenimine (Polysciences, Cat. # 23966-1)^80^. Medium was replaced with serum-free D-MEM the following day and collected 4 days later (5 days post-transfection). For purification, the medium was cleared of debris using a 0.45-µm filter (Corning, Part # 431220), loaded in two rounds onto a 100 kDa MW cutoff spin column (Amicon-Ultra-15, Merck Millipore, UFC910096) pre-treated with 1% BSA, and centrifuged at 3000g for 20 minutes. Concentrate was washed twice with Dulbecco’s PBS (DPBS, Gibco, cat. no. 14190-250) including 0.001% (vol/vol) Pluronic F-68 nonionic surfactant (Gibco, cat. no. 24040-032) at 3000g for 20 minutes yielding 200-300 μL of viral concentrate. This minimally purified solution was used for non-titered systemic delivery in mice through retro-orbital injections.

#### High-titer AAV production with purification

Recombinant adeno-associated viruses (AAVs) were produced following a previously described protocol^80^. HEK293T cells were maintained in Dulbecco’s Modified Eagle’s Medium (DMEM; Gibco) supplemented with 10% fetal bovine serum (FBS) and 1% penicillin–streptomycin (P/S) at 37 °C in a humidified atmosphere containing 5% CO₂. When cells reached approximately 80–90% confluency, the triple-plasmid transfection method was performed using PEI MAX® transfection reagent (Kyfora Bio). The gene of interest (GOI), pCAP (encoding Rep and Cap proteins), and pHelper (encoding adenoviral E4, E2A, and VA genes) plasmids (Addgene, #112865, #81070) were co-transfected at a ratio of 1:4:2. 24 hrs post transfection, the medium was replaced with DMEM containing 5% FBS and 1% P/S, and after 72 h trasnfection, both cells and medium were harvested. AAVs were purified by iodixanol gradient ultracentrifugation using a Type 70 Ti fixed-angle titanium rotor (Beckman-Coulter, Cat. No. 337922), followed by concentration and buffer exchange into Dulbecco’s phosphate-buffered saline (DPBS) containing 0.001% Pluronic F-68 (Gibco) using Ultracel centrifugal filters (Millipore) with a 100 kDa molecular-weight cutoff. Physical vector genome titers (vg/mL) were determined by droplet digital PCR (ddPCR) following the “ddPCR Titration of AAV Vectors” protocol (2) using a QX200 AutoDG Droplet Digital PCR System (Bio-Rad, Cat. No. 1864100). To ensure consistency, titration was performed using two primer–probe sets targeting ITR (Forward 5′-CGGCCTCAGTGAGCGA-3′, Reverse 5′-GGAACCCCTAGTGATGGAGTT-3′, Probe 5′-FAM-CACTCCCTCTCTGCGCGCTCG-3′) and WPRE (Forward 5′-GCTATGTGGATACGCTGCTTTA-3′, Reverse 5′-CCAGGATTTATACAAGGAGGAGAAA-3′, Probe 5′-HEX-TCATGCTATTGCTTCCCGTATGGCT-3′) regions.

### Stereotactic intraparenchymal AAV injection for local validation in mice and marmosets

#### Marmoset

For the final candidate screen and further validation of ubiquitous and projection-defined targeting with cjDAE8 in modified eAAV vector backbones, male marmosets (n=2, ∼3 years old) were implanted with a coordinate grid fixed to the skull and underwent T2-weighted structural MRI imaging for precisely targeted AAV injections to the ventral midbrain. To test the specificity of cjDAE4 and cjDAE8 in the original eAAV vector, one marmoset (20-160, aged 2.98 years) received injections of AAV1-iWPRE-cjDAE8-cm2-dTomato-WPRE-mBGHpA and AAV1-iWPRE-cjDAE4-cm2-EGFP-WPRE-mBGHpA (1:1 mix, 1E+12 vg/ml each, 1 μl/site at 100 nl/min) in the SNc and VTA. To test the specificity of cjDAE8 in the improved pAAV backbone and the ability to target projection-defined subpopulations of DA neurons through retrograde labeling, one male marmoset (23-116, aged 2.32 years) received stereotactic injections using the same targeting method. To assess the specificity of the improved backbone, the SNc was unilaterally injected at a lateral and medial site with AAV1-2xCHS4-cjDAE8-cm2-dTomato-WPRE-mHGHpA (1E+12 vg/ml, 1 μl/site at 100 nl/min). To assess selective labeling in projection-defined DA neurons, the caudate and putamen were injected with AAV2-retro-2xCHS4-cjDAE8-cm2-EGFP-WPRE-mHGHpA (5E+12 vg/ml, 2 μl/site at 200 nl/min). After 57 and 59 weeks of viral incubation, respectively, marmosets were euthanized and underwent transcardial perfusion in accordance with MIT CAC protocol as described previously. Brains were post-fixed overnight in 4% PFA 4°C, then serially sectioned by vibratome (100-μm thickness) in either the coronal or sagittal plane for immunofluorescence staining.

Briefly, midbrain serial sections underwent permeabilization with 2% PBST and then incubation for one hour at room temperature in blocking buffer solution containing 10% Blocking One buffer (Nacalai, Cat. # 03953-95) in 0.3% PBST. Blocked tissue sections were then incubated with mouse anti-TH IgG1 (1:2000, Immunostar, Cat. # 22941), chicken anti-GFP (1:2000, abcam, Cat. # ab13970), rabbit anti-RFP (1:1000, Rockland, Cat. # 600-401-379) primary antibodies in blocking buffer solution for 18-24 hours at 4°C. Next, tissue sections were incubated with goat anti-mouse Alexa-647 (1:500, Thermo Fisher Cat. # A21236), goat anti-rabbit Alexa-555 (1:500, Thermo Fisher Cat. # A21429), donkey anti-chicken Alexa-488 (1:1000, Thermo Fisher, Cat. # A78948), and DAPI (1:1000) to counterstain cell nuclei. To assess non-amplified endogenous reporter expression, adjacent serial sections were stained using the same protocol, but without anti-RFP and anti-GFP antibodies. To quench lipofuscin autofluorescence, the stained sections were treated for 20 minutes with TrueBlackPlus (1:40 in 1X PBS, Biotium, Cat. # 23014).

The specificity of endogenous versus antibody-amplified labeling were assessed using all stained coronal or sagittal sections (sampled every 100-μm for amplified, every 300-μm for endogenous) acquired with a FLUOVIEW FV3000 confocal microscope (10X objective, Olympus). A multi-tile Z-stack (40-μm, 5-μm step size) was acquired for each midbrain ROI. Maximum-intensity projections of these ROIs were used for subsequent automated analysis in custom pipelines using StrataQuest software. Whole-slide images were acquired by epi-fluorescent imaging using a VS200 microscope (10X objective, Olympus).

#### Mouse

To validate the specificity of cjDAE8 after direct local injection, 4-month-old male (n=2) and female (n=1) mice received stereotactic AAV injections under isoflurane anesthesia. AAV expressing 2xCHS4-cjDAE8-cm2-dTomato-WPRE-mHGHpA (AAV2/1, Boston Children’s Hospital Viral Core) was injected bilaterally (300 nL/site at 50nL/min, 5E+11 vg/mL titer) in either the VTA or SNc of each hemisphere at the following coordinates: SNc (AP: −3.40 mm, ML: +1.30 mm, DV: −4.20 mm from bregma) and VTA (AP: −3.50 mm, ML: −0.50 mm, DV: - 4.30 mm). Following 28 days of incubation, mice were euthanized with a fatal overdose of isoflurane anesthesia and underwent transcardial perfusion with ice-cold 1X PBS, followed by 4% PFA in 1X PBS. Brains were post-fixed overnight in 4% PFA, then serially sectioned coronally by vibratome (80-μm thickness) for immunofluorescence staining.

Briefly, midbrain serial sections underwent permeabilization with 2% PBST and then incubation for one hour at room temperature in blocking buffer solution containing 10% Blocking One buffer (Nacalai, Cat. # 03953-95) in 0.3% PBST. Blocked tissue sections were then incubated with mouse anti-TH IgG1 (1:2000, Immunostar, Cat. # 22941) and rabbit anti-RFP (1:1000, Rockland, Cat. # 600-401-379) primary antibodies in blocking buffer solution for 18-24 hours at 4°C. Next, tissue sections were incubated with goat anti-mouse IgG1 Alexa-488 (1:500, Thermo Fisher Cat. # A21121), goat anti-rabbit Alexa-647 (1:500, Thermo Fisher Cat. # A32733), and DAPI (1:1000) to counterstain cell nuclei.

The specificity of endogenous versus antibody-amplified labeling were assessed using three representative coronal sections per mouse (lateral, middle, and medial injection spread) acquired with a FLUOVIEW FV3000 confocal microscope (10X objective, Olympus). A multi-tile Z-stack (30-μm, 5-μm step size) was acquired for each midbrain ROI. Maximum-intensity projections of these ROIs were used for subsequent automated analysis in custom pipelines using StrataQuest software. Whole-slide images were acquired by epi-fluorescent imaging using a VS200 microscope (10X objective, Olympus).

### Specificity and efficiency of systemic DAE8 AAV administration in the mouse and marmoset

#### Retro-orbital AAV injection for mouse systemic validation

To validate the specificity and efficiency of cjDAE8 after systemic injection, 2-month-old male (n=2/dose) and female (n=2/dose) mice received retro-orbital AAV injections under isoflurane anesthesia at either a 5E+10 vg or 5E+11 vg dose in 1X PBS. After 28 days of incubation, mice were euthanized and perfused transcardially with ice-cold 1X PBS, followed by ice-cold 4% paraformaldehyde (PFA) fixative in 1X PBS. Brains were post-fixed overnight in 4% PFA at 4°C, then serially sectioned by vibratome (sagittal, 80-μm thickness) for immunofluorescence staining.

Briefly, serial sections underwent permeabilization with 2% PBST and then incubation for one hour at room temperature in blocking buffer solution containing 10% Blocking One buffer (Nacalai, Cat. # 03953-95) in 0.3% PBST. Blocked tissue sections were then incubated with mouse anti-TH IgG1 (1:2000, Immunostar, Cat. # 22941) and rabbit anti-RFP (1:1000, Rockland, Cat. # 600-401-379) primary antibodies in blocking buffer solution for 18-24 hours at 4°C. Next, tissue sections were incubated with goat anti-mouse IgG1 Alexa-488 (1:500, Thermo Fisher Cat. # A21121), goat anti-rabbit Alexa-647 (1:500, Thermo Fisher Cat. # A32733), and DAPI (1:1000) to counterstain cell nuclei.

The specificity and efficiency of endogenous versus antibody-amplified labeling were assessed with representative sagittal sections (lateral, middle, and medial) and acquired with a FLUOVIEW FV3000 confocal microscope (10X objective, Olympus). A multi-tile Z-stack (30-μm, 5-μm step size) was acquired for a generous midbrain ROI in each section. Maximum-intensity projections of these ROIs were used for subsequent analysis in custom pipelines with StrataQuest software. Identical fluorescence thresholds were used for positive nuclei detection across channels regardless of dose. Whole-slide images were acquired by epi-fluorescent imaging using a VS200 microscope (10X objective, Olympus).

#### Tail-vein AAV injection for marmoset systemic validation

To validate the specificity and efficiency of cjDAE8 after systemic delivery, we packaged the modified construct in AAV-BI103, an optimized AAV variant featuring efficient blood-brain-barrier transport and widespread CNS transduction. One female marmoset (22-132, aged 2.6 years) weighing 300 g was anesthetized and injected in the tail vein with AAV-BI103-2xCHS4-cjDAE8-cm2-dTomato-6xmiR-WPRE3-mHGHpA (8E+13 vg/kg dose). After 7.6 weeks of viral incubation, the marmoset was euthanized as described previously and underwent transcardial perfusion with ice-cold 1X PBS followed by ice-cold 4% paraformaldehyde (PFA) in 1X PBS. The brain was post-fixed overnight in 4% PFA, then one hemisphere was serially sectioned sagittally by vibratome (100-μm thickness) for immunofluorescence staining. The other hemisphere was further post-fixed with SHIELD buffer (LifeCanvas, Cat. # GE38) for whole-tissue clearing with immunofluorescent staining.

For traditional immunofluorescence in serial sections, slices were permeabilized with 2% PBST and then incubated for one hour at room temperature in blocking buffer solution containing 10% Blocking One buffer (Nacalai, Cat. # 03953-95) in 0.3% PBST. Blocked tissue sections were then incubated with mouse anti-TH IgG1 (1:2000, Immunostar, Cat. # 22941) and rabbit anti-RFP (1:1000, Rockland, Cat. # 600-401-379) primary antibodies in blocking buffer solution for 18-24 hours at 4°C. Next, tissue sections were incubated with goat anti-mouse IgG1 Alexa-488 (1:500, Thermo Fisher Cat. # A21121), goat anti-rabbit Alexa-647 (1:500, Thermo Fisher Cat. # A32733), and DAPI (1:1000) to counterstain cell nuclei. To quench lipofuscin autofluorescence, the stained sections were treated for 20 minutes with TrueBlackPlus (1:40 in 1X PBS, Biotium, Cat. # 23014).

We characterized the specificity of endogenous versus antibody-amplified labeling using images of all stained sagittal sections (sampled every 100-μm) acquired with a FLUOVIEW FV3000 confocal microscope (10X objective, Olympus). A multi-tile Z-stack (40-μm, 5-μm step size) was acquired for each midbrain ROI. Maximum-intensity projections of these ROIs were used for subsequent automated analysis in custom pipelines using StrataQuest software. Whole-slide images were acquired by epi-fluorescent imaging using a BX61WI microscope (4X objective, Olympus).

All clearing, immunofluorescent staining, and light-sheet imaging protocols were performed by LifeCanvas Technologies, CRO. After SHIELD post-fixation, the whole marmoset hemisphere was cleared for 6 weeks with SmartBatch+. All staining protocols were performed following the SmartBatch+ Radiant Protocol. Cleared tissue was incubated with 60 µg rabbit anti-RFP (Rockland, Cat. # 600-401-379), 72 µg mouse anti-TH (BioLegend, Cat. # 818001). Next, the tissue was incubated sequentially with donkey anti-rabbit Alexa-555 (Invitrogen, Cat. # A32794) and donkey anti-mouse SeTau Alexa-647 (LifeCanvas Technologies, Cat. # DkxMs-ST). The sample was index-matched with EasyIndex (RI=1.52). After first-pass imaging, tissue was re-labeled with 72 µg mouse anti-TH using simultaneous Fabs (Jackson) due to the low penetrance of TH labeling in lateral tissue regions. 3D images were acquired by light-sheet microscopy with SmartSPIM at 3.6X magnification (4 µm z-step, 1.8 µm xy pixel size). Raw TIF images were processed using Imaris software for 3D reconstruction, volumetric re-slicing, and video generation.

### Semi-automated cell quantification and colocalization analysis with StrataQuest

Acquired ROI confocal images of serial sections across histology datasets were analyzed using custom pipelines in StrataQuest software with manual validation of fluorescent intensity thresholds for a subset of images in each pipeline. Briefly, gray virtual channel composites (overlays of 488, 555, and 647 channels) were dilated and eroded to enrich fluorescence from cell bodies over neurites. Next, we trained a random forest model on a representative subset of images per dataset to classify cell bodies. The model was trained on the eroded and median-filtered gray virtual channel composites using a pixel size of 3 (mouse) or 5 (marmoset) and smoothing (size=1). Upon sufficient training, the model classified cell bodies across all images. We then masked the original composites with this output for nuclei segmentation of cell bodies selectively over neurites. Positive detections were set by thresholding fluorescence intensity of each channel across nuclei to achieve consensus with manual cell counts.

### Fiber photometry recordings of CS and DLS DA terminals during head-fixed locomotion and air puff stimuli

Female mice (n=4, 2 months old) received bilateral injections in the SNc (AP: −3.40 mm, ML: +/-1.30 mm, DV: −4.10 mm) with AAV9-iWPRE-cjDAE8-cm2-jGCaMP6f-WPRE-BC-bGHpA (300 nL/site, 8E+12 vg/mL) and then implanted optic fiber (400-µm core, 0.5NA, RWD) cannulas in the CS (AP: +0.60 mm, ML: -/+1.80 mm, DV: −3.40 mm) and DLS (AP: +0.60 mm. ML: +/-2.10 mm, DV: −2.90 mm) in opposite hemispheres. Left versus right DLS and CS implantation was counterbalanced across mice to mitigate the possibility of confounding hemisphere-specific movement signals. After 3-4 weeks of viral incubation, mice were habituated to the head-fixed treadmill apparatus (LabeoTech, Cat. # 38202) and patch cables over two sessions, with the first session lasting 10 minutes and the second session lasting 30 minutes. To assess whether DA terminal activity in the CS and DLS differentially correlate with movement, synchronized fiber photometry and treadmill (distance, velocity) recordings were acquired at 25 Hz and 100 Hz, respectively, over two 30-minute sessions separated by 5 days.

Fiber photometry data were acquired with equal power LEDs (410 nm, 470 nm; 30-40 µW power) using the R820 photometry system and acquisition software (RWD). For sessions recording movement velocity, the R820 sent triggered TTL pulses to the treadmill apparatus for synchronized locomotion data with the fluorescent signals. To assess whether DA terminal activity in the CS and DLS differentially responds to air puff stimuli, mice underwent two sessions receiving 9 or more trials of unanticipated air puffs delivered manually (variable ITI, 20-90 sec) to the rear fur while head-fixed on the treadmill.

In the RWD photometry analysis software, the raw fluorescence signals were smoothed (W = 5), baseline-corrected (β = 8), and motion-corrected using the isosbestic (410 nm) signal. Next, the DF/F and Z-score data were computed using a 60-second time window. The average Pearson cross-correlation between velocity/acceleration and Z-scored CS/DLS DA activity was calculated and plotted using custom Python scripts. First, the loaded velocity (cm/s) and calculated acceleration (cm/s^2^) data were resampled and aligned to match the fluorescence timestamps and smoothed with a Gaussian filter (σ=5). Z-scored CS and DLS fluorescence traces were masked during periods of movement in which speed and acceleration exceeded 5 cm/s and 50 cm/s^2^, respectively. For each mouse and region, we computed Pearson cross-correlations between fluorescence and (i) speed and (ii) acceleration across lags ±5 s during these movement periods. Per-mouse summary values were generated for velocity at τ=0 s and acceleration averaged over −1<τ<0 s versus 0<τ<+1 s. These individual mean Pearson r values underwent a two-way repeated-measures ANOVA (Region × Period) with Šídák multiple-comparisons (Prism), yielding the interaction and pairwise contrasts reported. Mean correlations were averaged across sessions per mouse, which were averaged across all mice to generate the final group-level data and plots.

For the analysis of air-puff triggered DA terminal responses, Z-scored CS and DLS fluorescence were aligned to air puff onset (t=0) across trials and averaged across sessions for each mouse. The mean triggered-responses for CS and DLS were then averaged across mice for the final group-level data (mean ± SEM). Area under the curve (AUC) analysis was performed on the mean Z-score trace for each animal over fixed time windows before (pre, −0.75–0 s) and after (post, 0–0.75 s) air puff onset. Per-mouse AUCs were averaged across trials to give one value per Region×Time cell. Group statistics used a two-way repeated-measures ANOVA (Region × Time) with Bonferroni-corrected post-hoc comparisons.

### VTA-DA self-stimulation with optogenetics

For optogenetic self-stimulation experiments, male (n=4) and female (n=6) mice aged 2-4 months old received bilateral AAV injections in the VTA at the following coordinates (relative to bregma): AP −2.90 to −3.10 mm, ML +/-0.55 to 0.60 mm, DV −4.55 mm. A 200-µm core optic fiber (0.39 NA, RWD) was slowly implanted unilaterally 150-µm above the injection site and cemented to the skull. Mice were injected with 300 nL (at 50 nl/min) of either AAV1-2xCHS4-cjDAE8-cm2-ChR2(H134R)-EYFP-WPRE-mHGHpA (n=6) or the control AAV1-2xCHS4-cjDAE8-cm2-dTomato-WPRE-mHGHpA (1E+12 vg/mL, n=4). Mice injected with the ChR2(H134R) construct received one of the following titers: 1E+13 vg/mL (n=1), 5E+12 vg/mL (n=1), 2E+12 vg/mL (n=2), or 1E+12 vg/mL (n=2).

After at least 14 days of viral incubation, mice underwent three days optogenetic self-stimulation training (30-min duration, 180 maximum stimulations). Each session, mice were placed in an operant chamber (Part # ENV-307A-CT, Med Associates) equipped with two nose-poke ports (Part # ENV-313M, Med Associates) and LED lights (Part # ENV-321DM, Med Associates). Each trial began with the illumination of the LED cue light. A single nose poke at the active port during cue light illumination triggered a 2-s photostimulation (10-ms pulse width, 20 Hz, 4.5-5 mW, 20% duty cycle) of VTA-DA neurons along with a 2-s illumination of the port. At the end of stimulation, cue and nose-poke LED lights turned off for an 8-s inter-trial interval (ITI) during which active pokes were not reinforced by photostimulation.

After training, mice were euthanized with a fatal overdose of isoflurane anesthesia and underwent transcardial perfusion with ice-cold 1X PBS, followed by 4% PFA in 1X PBS. Brains were post-fixed overnight in 4% PFA, then serially sectioned coronally by vibratome (80-μm thickness) for immunofluorescence staining. Briefly, midbrain serial sections underwent permeabilization with 2% PBST and then incubation for one hour at room temperature in blocking buffer solution containing 10% Blocking One buffer (Nacalai, Cat. # 03953-95) in 0.3% PBST. Blocked tissue sections were then incubated with mouse anti-TH IgG1 (1:2000, Immunostar, Cat. # 22941) in blocking buffer solution for 18-24 hours at 4°C. Next, tissue sections were incubated with goat anti-mouse 647 (1:500, Thermo Fisher, Cat. # A21236) and DAPI (1:1000) to counterstain cell nuclei. Representative coronal sections per mouse (lateral, middle, and medial to injection/implantation site) were acquired by confocal microscopy with the FLUOVIEW FV3000 (Olympus) or LSM700 (Zeiss).

### Data availability

To promote transparency and support the research community, all histology datasets will be hosted for at least a year via the Neuroglancer viewer and will be permanently available to download through Zenodo [<u>mouse datasets</u> DOI: 10.5281/zenodo.17645143; <u>marmoset datasets</u> DOI: 10.5281/zenodo.17644916]. These resources include: (i) high-resolution confocal images of midbrain regions used to quantify cjDAE8-AAV performance and corresponding StrataQuest analysis pipelines, and (ii) whole-slide epifluorescent images. Additionally, 3D light-sheet imaging data from a cleared marmoset hemisphere—following intravenous cjDAE8-AAV delivery and immunofluorescent staining for RFP and TH—are hosted on neuroglancer to enable comprehensive, whole-brain evaluation of tool performance. Neuroglancer dataset descriptions and access links are <u>available on GitHub</u> if there are any issues with the direct hyperlinks referenced below. Along with histology data, all raw and processed data for fiber photometry and optogenetics experiments will be available for long-term access through Zenodo [<u>functional</u> <u>validation datasets</u> DOI: 10.5281/zenodo.17645091].

#### Marmoset Neuroglancer Data

To view confocal images of midbrain ROIs after local injection in marmoset, click <u>here</u> for the native fluorescence data and click <u>here</u> for the antibody-amplified fluorescence data. Furthermore, to view whole-slide epifluorescent images of the marmoset local injection data, click <u>here</u> for the native fluorescence dataset and click <u>here</u> for the antibody-amplified fluorescence dataset. To view the 3D-cleared marmoset hemisphere with anti-RFP and anti-TH immunostaining after systemic delivery, click <u>here</u>. To view confocal images of midbrain ROIs in serial sections with anti-RFP and anti-TH immunostaining, using the other hemisphere after systemic delivery, click <u>here</u>.

#### Mouse Neuroglancer Data

To view whole-slide epi-fluorescent images of low dose (5E+10 vg) systemic delivery with anti-RFP and anti-TH immunostaining, where each slide is a series from a different mouse, click <u>here</u>. To view whole-slide epi-fluorescent images of high dose (5E+11 vg) systemic delivery with anti-RFP and anti-TH immunostaining, where each slide contains the series of a different mouse, click <u>here</u>. Finally, to view whole-slide epi-fluorescent images of mouse local injection with anti-RFP and anti-TH immunostaining, where each slide contains the series of a different mouse, click <u>here</u>.

## Code availability

Custom pipelines Jupyter notebooks used for fiber photometry analyses and 2D neuroglancer dataset generation are available to download on GitHub.

## Acknowledgements

This research was supported by the NIH BRAIN Initiative Grant UG3 MH126869 (OSP#6947909). K.A.C. was supported by the BCS Alder (1972) Graduate Student Fellowship and the NSF Graduate Research Fellowship.

We thank Kirsten Levandowski, Qiangge Zhang, and Yefei Chen for performing various marmoset procedures (intravenous injections, euthanasia, tissue collection), as well as Will Menegas for feedback on figure formatting. We also thank Brian Nguyen and his colleagues at LifeCanvas Technologies for performing the marmoset tissue clearing experiment. We deeply thank Russell Ulbrich and his colleagues for training K.A.C. on StrataQuest and their technical assistance with developing analysis pipelines.

Finally, we express our deepest gratitude and respect for all the animals euthanized for this work. We also want to recognize the invaluable contributions of marmosets Ivy (19-184), Florian (20-160), Swan (23-116), and Dove (22-132) to the development of this technology.

## Author contributions

K.A.C. designed and oversaw research methodology for the candidate screening, enhancer-backbone development, and *in vivo* characterization of cjDAE-AAVs under the supervision of G.F., F.W., M.W., and F.K.; F.K., Y.W. sequenced and analyzed marmoset multiomics data; R.R. analyzed conservation of analogous enhancer sequences and regulatory elements across species; K.A.C. designed and cloned all pAAV constructs; C.S.C and M.W. packaged all AAVs with minimal purification approach; I.K. packaged all high-titer purified AAVs; K.A.C. performed initial cjDAE candidate screening in mouse; K.A.C., C.S.C., and M.W. designed and executed enhancer-backbone optimization experiments (data not shown); K.A.C performed all systemic and local injections in mice, R.C. and M.W. performed all marmoset local injection procedures; K.A.C. designed and executed all histology analyses. K.A.C. designed, performed, and analyzed data for fiber photometry experiments; K.C. and K.A.C. designed, performed, and analyzed data for optogenetics experiments; K.A.C. wrote the manuscript, and K.C., M.W., Y.W., and I.K. wrote methods subsections; all authors edited and approved.

## Ethics declarations

All animal procedures were performed in accordance with MIT CAC protocols. No subjects were excluded from any of the analyses. For SNc fiber photometry experiments, researchers were blinded to CS versus DLS recording sites during data acquisition and processing. For VTA optogenetics experiments, researchers were not blinded.

## Reporting summary

A reporting summary will be provided as a separate file upon submission.

---

**Extended Data Figure 1.**
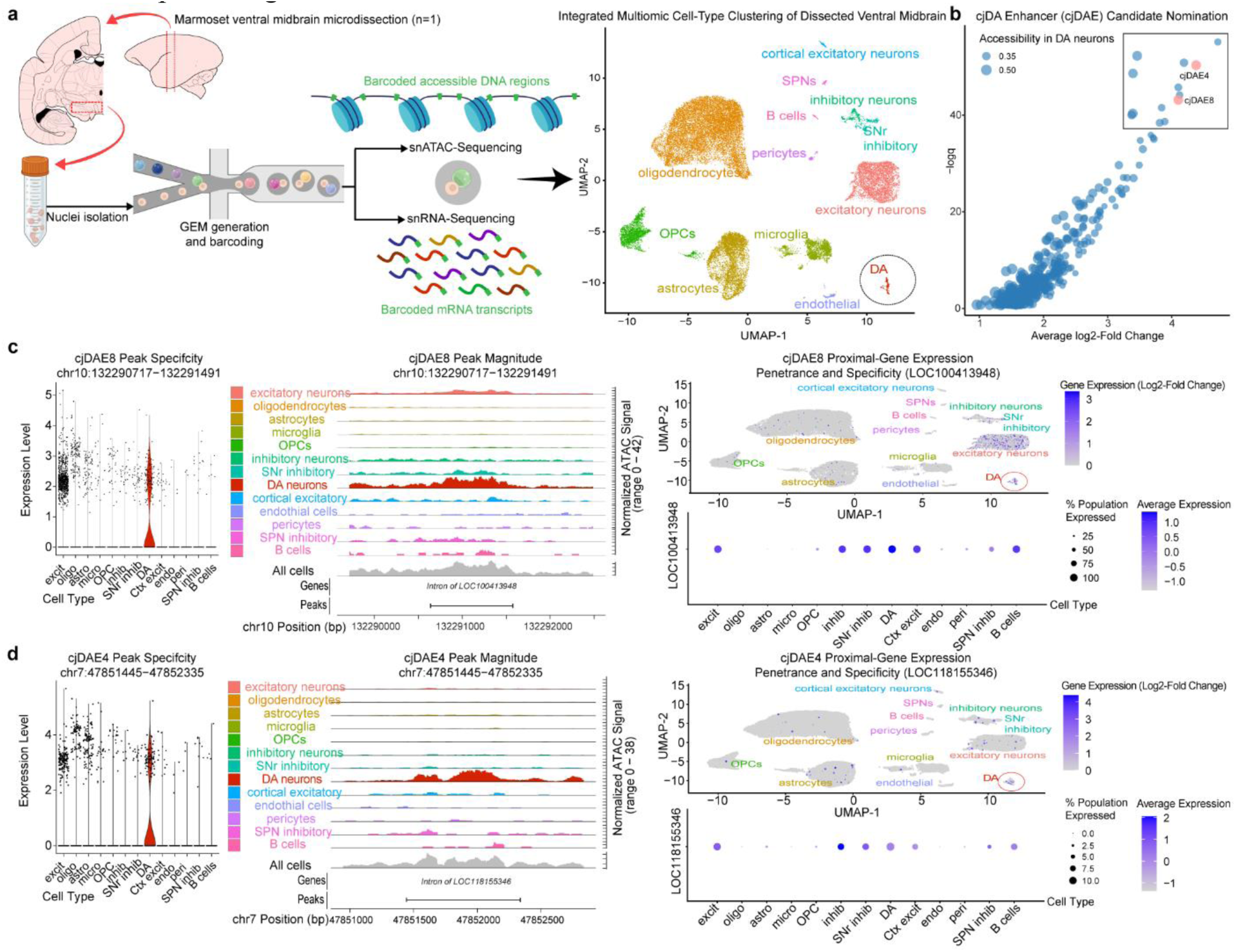
Nomination of marmoset DA-neuron enhancer (cjDAE) candidates. **a,** Dual snRNA- and snATAC-sequencing of unsorted nuclei was employed on freshly dissected marmoset midbrain tissue for integrated multiomic analysis of differentially accessible chromatin peaks across transcriptionally defined cell populations (2x reactions, 32,070 total high-quality nuclei). Candidate enhancer peaks were nominated by identifying chromatin regions with penetrant accessibility and specificity to the DA-neuron cluster (258 total nuclei, circled in black) compared to all others. Schematics made in BioRender. **b,** Peaks were further filtered to 401 candidates using thresholds for penetrance of accessibility (>30% in DA cluster, <5% in non-DA) and normalized magnitude of peak expression ( ≥1.5 log2-fold change). The final 12 candidates, such as cjDAE4 (**c**) and cjDAE8 (**d**), were selected from the ranked filtered list after further considering the genome context (enrichment of DA-related genes most proximal to the enhancer in linear-sequence distance).

**Extended Data Figure 2.**
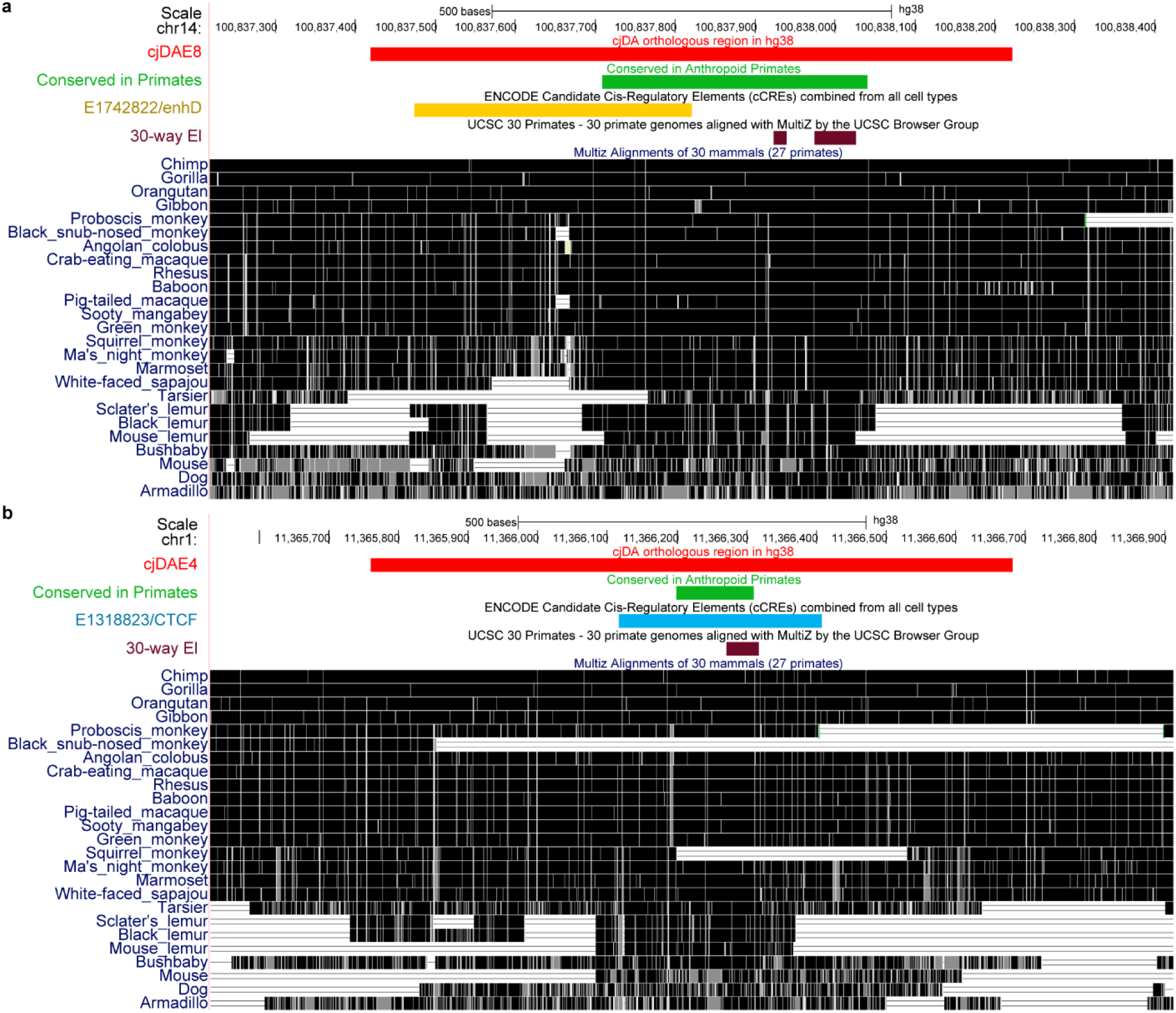
Evolutionary conservation of cjDAE8 and cjDAE4 sequences, as well as cis-regulatory element signals of respective human orthologs. **a,** Human orthologous region of cjDAE8 (track labeled “cjDA orthologous region in hg38”; red region). Multiple sequence alignment shows overall conservation in primates with strong conservation in great apes and old-world monkeys, and very weak conservation in non-primate mammals (“Multiz Alignments of 30 mammals (27 primates)”; darker shades indicate high sequence similarity with human)^81^. The cjDAE8 human ortholog contains a subregion highly conserved in human, orangutan, rhesus macaque and marmoset (“Anthropoid Primate Conservation”; green region)^82^, which further contains two short regions that are highly conserved across primates and non-primate mammals (“UCSC 30 Primates – 30 primate genomes aligned with MultiZ by the UCSC Browser Group”; maroon region)^83^. A candidate cis-regulatory element identified by Encode to have a distal enhancer-like signature lies on the left half of the cjDAE8 human ortholog (“ENCODE Candidate Cis-Reglatory Elements (cCREs) combined from all cell types”; yellow region)^84^. **b**, Human orthologous region of cjDAE4 (“cjDA orthologous region in hg38”; red). Multiple sequence alignment shows strong conservation in great apes and weaker conservation in Old World and New World monkeys (“Multiz Alignments of 30 mammals (27 primates)”). The human ortholog of cjDAE4 contains a subregion highly conserved in anthropoid primates (“Anthropoid Primate Conservation”; green), which contains a shorter region that is highly conserved across mammals (“UCSC 30 Primates – 30 primate genomes aligned with MultiZ by the UCSC Browser Group”; maroon). The highly conserved regions in anthropoid primates (green) and mammals (maroon) lie in a candidate cis-regulatory element identified by Encode to have high DNAse and CTCF and low H3K4me3 and H3K27ac (“ENCODE Candidate Cis-Reglatory Elements (cCREs) combined from all cell types”; light blue region).

**Extended Data Figure 3.**
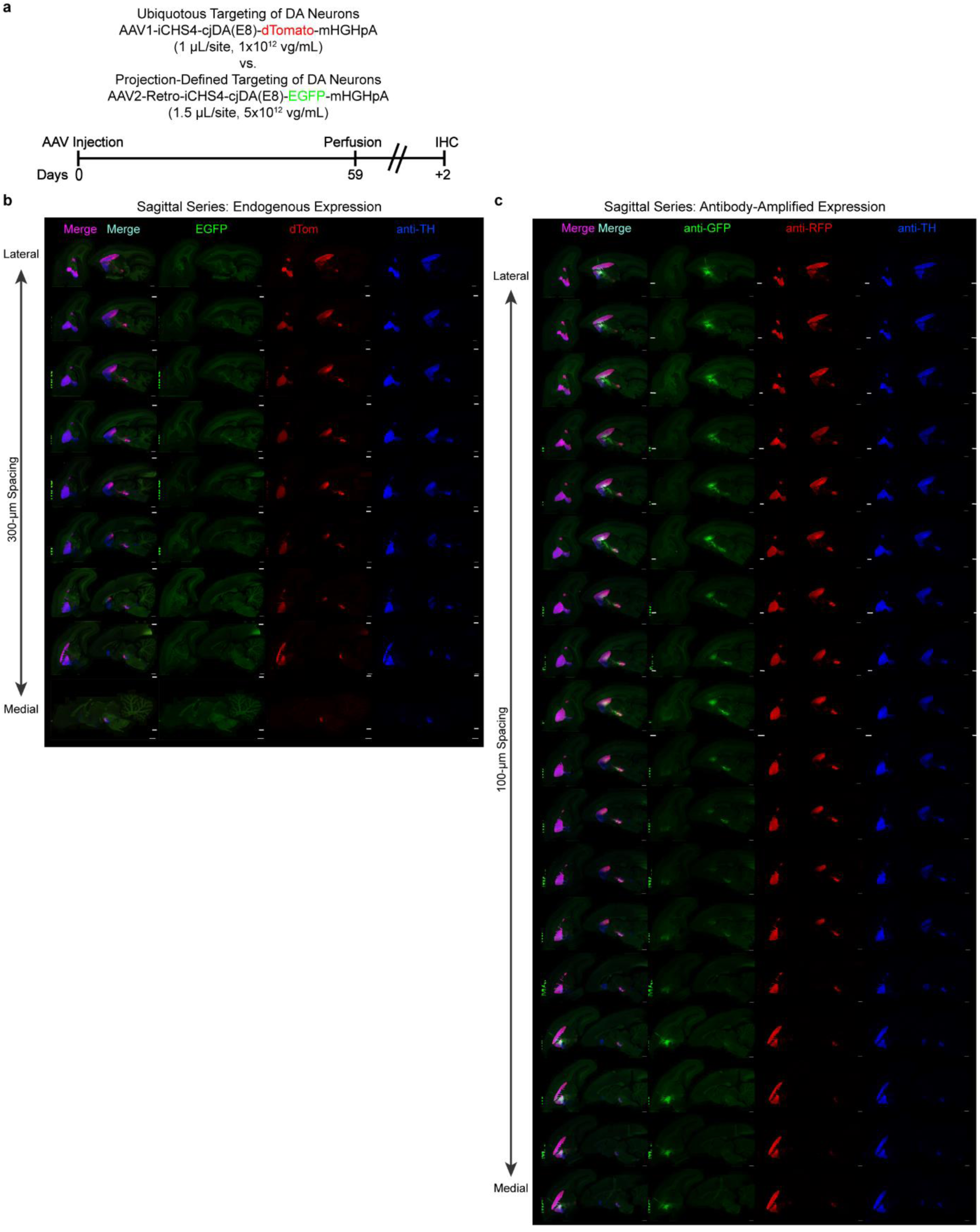
Strategies of locally delivering backbone-modified cjDAE8-AAVs in marmoset for ubiquitous or projection-defined targeting of DA neurons with unprecedented specificity. **a,** Revisited local injection strategy for ubiquitous and projection-defined targeting of DA neurons in the marmoset. Schematics made in BioRender. **b,** Serial sagittal epifluorescence images spanning the full dataset for the endogenous (non-amplified) reporter condition, showing robust midbrain-focused cjDAE8-dTom and cjDAE8-EGFP labeling patterns across sections. Scale bar, 2 mm. **c,** Serial sagittal epifluorescence images sampling the entire hemisphere the antibody-amplified reporter condition, revealing increased signal and fine terminal fields while preserving DA-neuron specificity achieved by the backbone-optimized cjDAE8 construct. Scale bar, 2 mm; channels shown as EGFP/dTom/anti-TH (endogenous) and anti-GFP/anti-RFP/anti-TH (amplified). Scale bar, 500 μm.

**Extended Data Figure 4.**
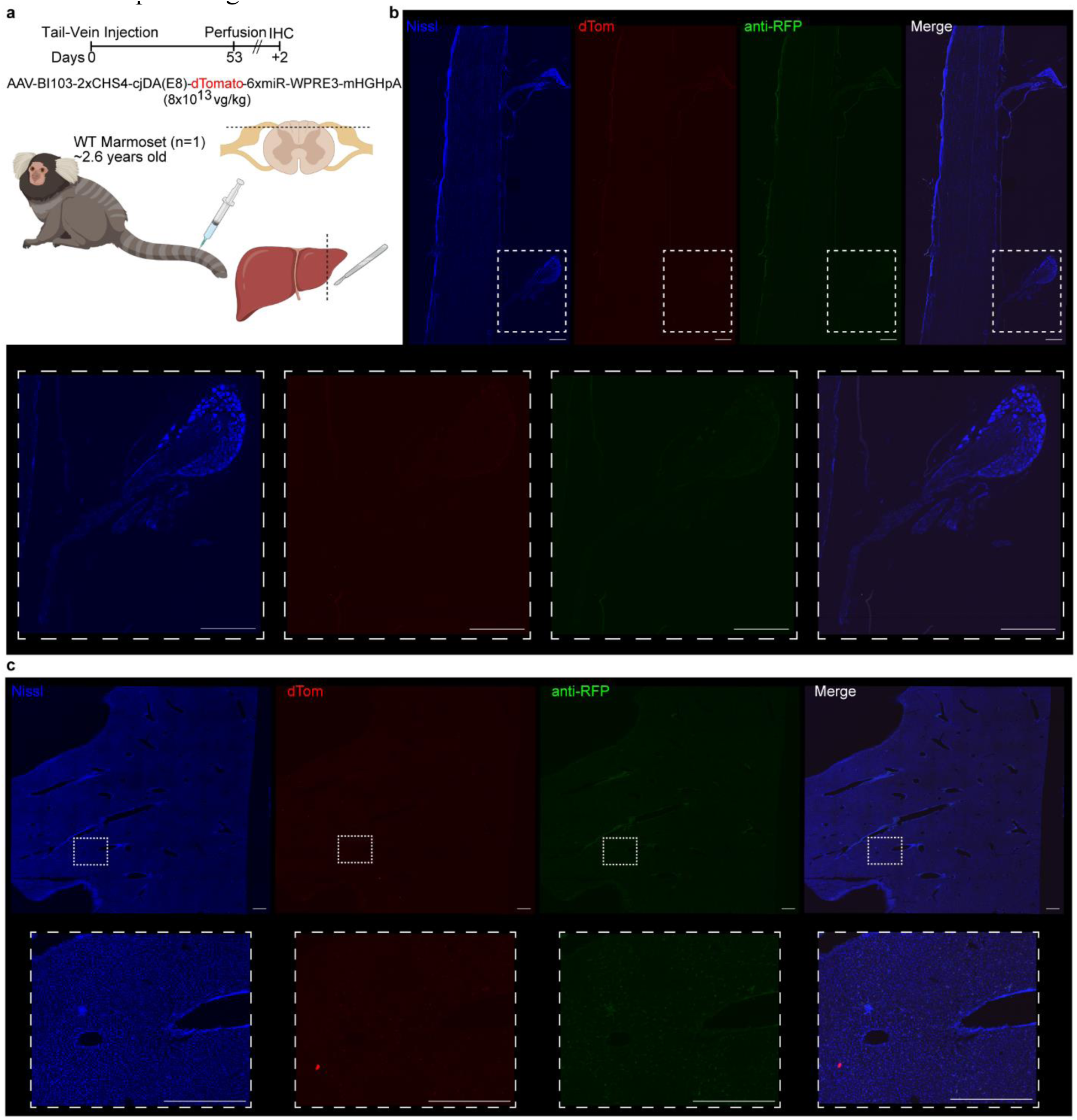
Transgene expression is undetectable in the DRG and liver after systemic delivery of cjDAE8-AAV in the marmoset. **a,** Revisited experimental timeline and strategy for systemic targeting of DA neurons in marmoset using AAV-BI103, with an additional schematic of peripheral tissues processed. **b,** Endogenous and amplified cjDAE8-dTomato expression was not apparent in sections of the DRG and spinal cord. Scale bar, 500 µm. **c,** Similarly, labeling with either was undetected in sections of the liver. Scale bar, 500 µm.

**Extended Data Figure 5.**
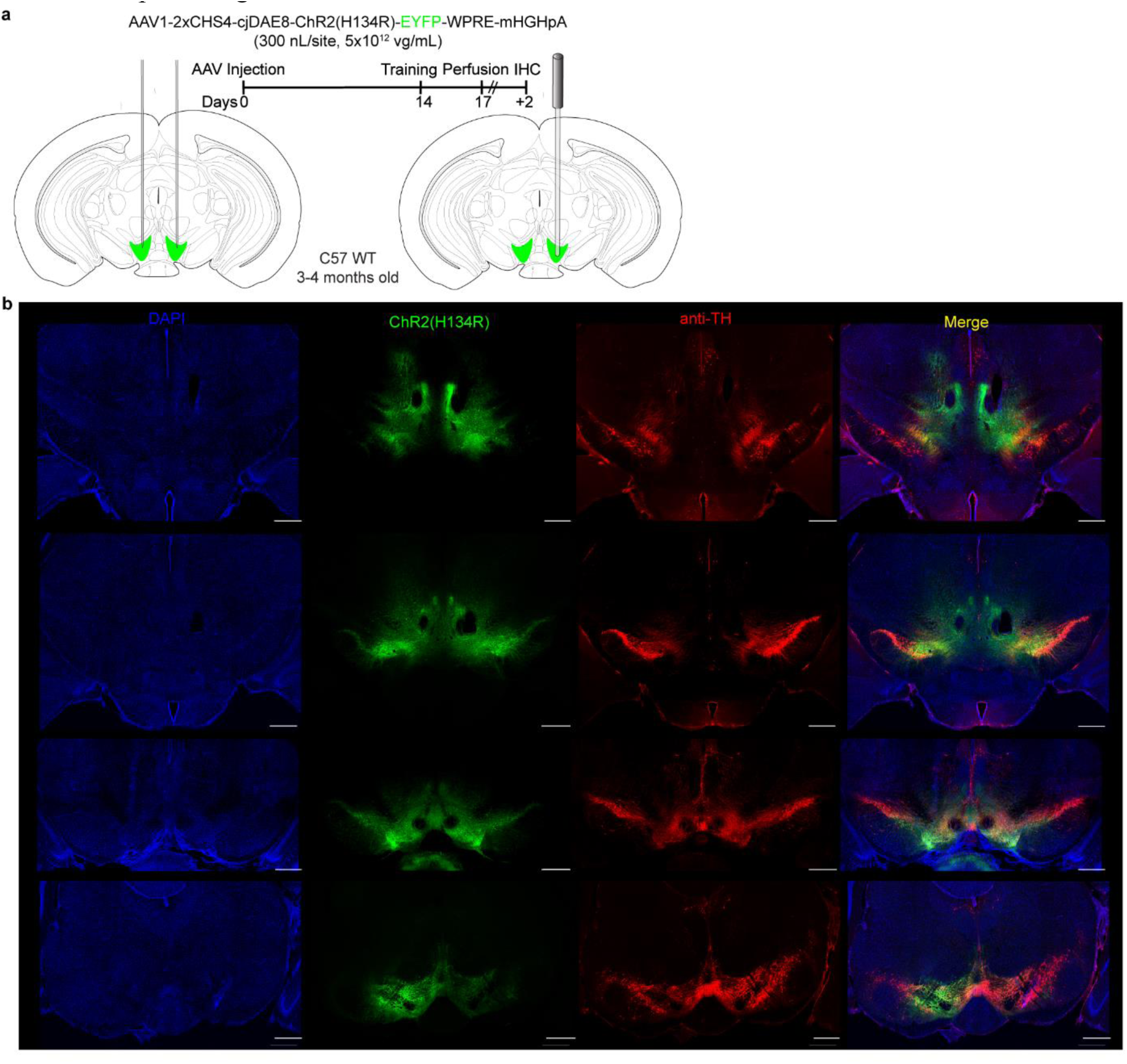
High-titer local injection of cjDAE8-ChR2(H134R)-EYFP increases off-target expression. **a,** Revisited experimental timeline and strategy for the VTA-DA self-stimulation experiment, using a 300-nl injection volume to test a range of titers. Mice receiving high titers (5E+12 or 1E+13 vg/ml) showed notably reduced labeling specificity. In this example, the mouse was injected with a 5E+12 vg/mL titer. **b,** Notable off-target EYFP labeling surrounding the injection site, with reduced TH-staining levels in sections with strong labeling. Scale bar, 500 µm.

## Supplementary information

Supplementary Videos 1-2 contain movies of the 3D cleared marmoset hemisphere with cjDAE8 labeling in the coronal and sagittal plane.

